# Identification of new proviral and antiviral factors through the study of the Dicer-2 interactome *in vivo* during viral infection in *Drosophila melanogaster*

**DOI:** 10.1101/2023.12.21.570062

**Authors:** Claire Rousseau, Émilie Lauret, Lauriane Kuhn, Johana Chicher, Philippe Hammann, Carine Meignin

## Abstract

RNA interference, which has a major role in the control of viral infection in insects, is initialized by the sensing of double stranded RNA (dsRNA) by the RNAse III enzyme Dicer-2. Although many *in vitro* studies have helped understand how Dicer-2 is able to discriminate between different dsRNA substrate termini, much less is known about how this translates to the *in vivo* recognition of viral dsRNA. Indeed, although Dicer-2 associates with several dsRNA-binding proteins (dsRBPs) that can modify its specificity for a substrate, it remains unknown how Dicer-2 is able to recognize the protected termini of viral dsRNAs. In order to study how the ribonucleoprotein network of Dicer-2 impacts antiviral immunity, we used an IP-MS approach to identify *in vivo* interactants of different versions of GFP::Dicer-2 in transgenic lines. We provide a global overview of the partners of Dicer-2 *in vivo*, and reveal how this interactome is modulated by different factors such as the viral infection and/or different point mutations inactivating the helicase or RNase III domains of GFP::Dicer-2. Our analysis uncovers several previously unknown Dicer-2 interactants associated with RNA granules (i.e. Me31B, Rump, eIF4E1 & Syp). Functional characterization of the candidates reveals pro- and antiviral factors in the context of the infection by the picorna-like DCV virus. In particular, the protein Rasputin has been identified as a novel antiviral candidate. The resources provided by this work can be used to gain a better understanding of the molecular complexes assembled around Dicer-2 in the context of antiviral RNAi and beyond.

## INTRODUCTION

In metazoans, virus-derived double-stranded RNAs (dsRNAs) allow the detection of a broad range of viruses by pattern recognition receptors (PRRs), leading to the activation of antiviral innate immunity (Rousseau & Meignin, 2020). Dicer proteins are dsRNA sensors with an endoribonuclease activity from the RNase III family, enabling the production of microRNAs (miRNAs) and small interfering RNAs (siRNAs) (Baldaccini & Pfeffer, 2021). *Drosophila melanogaster*, like other arthropods, encodes two Dicer proteins: Dicer-1 which is dedicated to miRNA processing, and Dicer-2 which is required for siRNA biogenesis (Lee *et al*, 2004). Insects largely rely on the siRNA pathway for antiviral defense through the detection of viral dsRNA by Dicer-2 (Galiana-Arnoux *et al*, 2006; van Rij *et al*, 2006; Kemp *et al*, 2013; Webster *et al*, 2015; de Faria *et al*, 2022; Wang *et al*, 2006). The virus-derived siRNAs (vsiRNAs) produced by Dicer-2 are then loaded onto the protein Argonaute2 (AGO2) to form the RNA-induced silencing complex (RISC), which targets and degrades the complementary RNA (Hammond *et al*, 2001; van Rij *et al*, 2006; Wang *et al*, 2006). Dicer-2 has also been proposed to participate in the regulation of the expression of antiviral genes, in addition to its function in RNAi (Deddouche *et al*, 2008; Göertz *et al*, 2019).

Dicer-2 is able to recognize and discriminate between two types of dsRNA termini and subsequently initiates two distinct types of cleavage mechanisms, called processive and distributive dicing (Welker *et al*, 2011; Cenik *et al*, 2011; Sinha *et al*, 2015, 2018). *In vitro*, blunt dsRNA promotes processive cleavage, whereby the helicase domain of Dicer-2 will bind the dsRNA termini and thread through the dsRNA molecule using ATP hydrolysis to produce multiple siRNA duplexes in one go. In contrast, dsRNA with a 3’overhang promotes distributive cleavage, whereby the 5′-monophosphate of the dsRNA substrate is anchored by the phosphate-binding pocket in the Dicer-2 Platform·PAZ domain (Kandasamy & Fukunaga, 2016; Kandasamy *et al*, 2017) and Dicer-2 dissociates after each high-fidelity cleavage in an ATP-independent manner (Donelick *et al*, 2020).

The helicase domain of all Dicer proteins is phylogenetically related to the retinoic acid-inducible gene-I (RIG-I)-like receptors RLRs (Deddouche *et al*, 2008; Baldaccini & Pfeffer, 2021). RLRs are cytosolic dsRNA sensors that induce an interferon response in vertebrates, suggesting that the helicase domain is a sensor of viral infection. Viral dsRNAs can be synthetized as an intermediate product of viral replication in RNA viruses or can originate from the convergent transcription of DNA viruses. In drosophila, all viruses tested so far are detected by Dicer-2 and induce the production of vsiRNAs (van Rij *et al*, 2006; Wang *et al*, 2006; Kemp *et al*, 2013; Mueller *et al*, 2010; de Faria *et al*, 2022). However, as the dicing mechanism relies on the recognition of the termini of the substrate, and viral dsRNAs produced during viral replication *in vivo* do not usually contain free ends, this raises the question of how Dicer-2 and the RLRs are able to sense viral infection. For example, the genome of the picorna-like *Drosophila C virus* (DCV, *Dicistroviridae*) is protected at the 5’end by a covalently bound protein called VPg and at the 3’end by a polyA tail.

To perform its function, Dicer-2 associates with several dsRNA-binding proteins (dsRBPs). It has been shown that the two accessory dsRBPs Loqs and R2D2 both bind the helicase domain of Dicer-2, consisting of 3 subdomains, Hel1, Hel2i and Hel2 (**Figure 1A**) (Lim *et al*, 2008; Nishida *et al*, 2013; Yamaguchi *et al*, 2022). More specifically, Loqs binds the Hel2 domain (Trettin *et al*, 2017) and R2D2 binds the Hel2i domain (Yamaguchi *et al*, 2022). Other dsRBPs (e.g. TRBP, PACT, PKR and ADAR) have also been shown to interact with the helicase domain of human Dicer (Ota *et al*, 2013; Wilson *et al*, 2015; Liu *et al*, 2018b; Montavon *et al*, 2021). The helicase domain therefore appears to be of utmost importance for the interaction of Dicer-2 with different regulatory proteins. Moreover, it has been shown that the helicase domain of human Dicer and drosophila Dicer-2 is important for antiviral response (Deddouche *et al*, 2008; Marques *et al*, 2013; Poirier *et al*, 2018; Donelick *et al*, 2020; Montavon *et al*, 2021).

**Figure 1:**
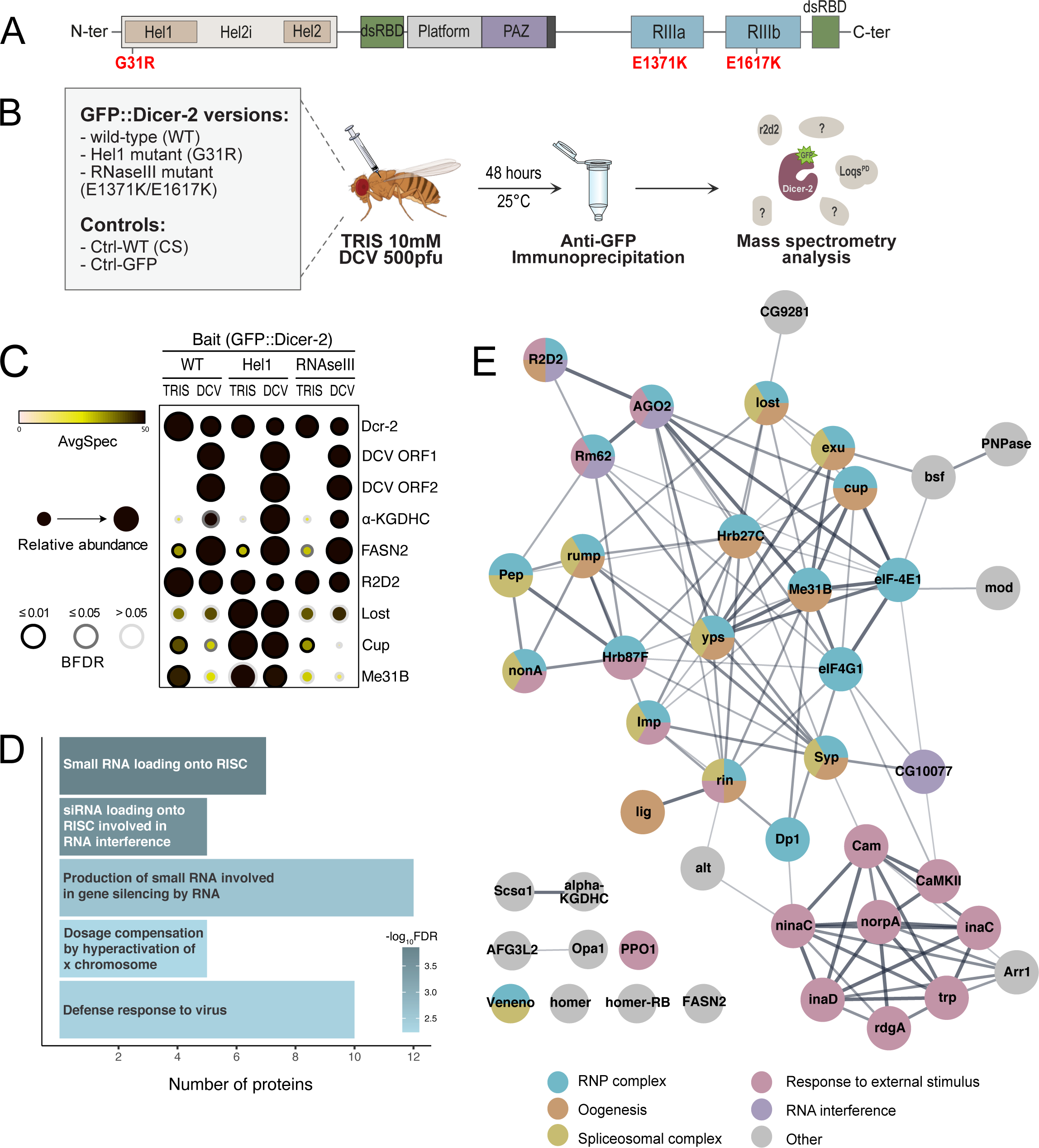
Global Dicer-2 interactome network during viral infection *in vivo*. **(A)** Schematic representation of Dicer-2 domain architecture. The Dicer-2 protein (1722 amino acids) is composed of a helicase domain (which contains the Hel1, Hel2i and Hel2 sub-domains), two double-strand RNA binding domains (dsRBD), a PAZ domain, and two ribonuclease III domains (RIIIa and RIIIb). G31R and E1371K/E1671K represent the point mutations of *dicer-2* used in this study. **(B)** Scheme illustrating the experimental strategy for IP & LC-MS/MS used to identify Dicer-2 partners *in vivo* in mock-infected (TRIS) and DCV-infected conditions. All genotypes used are presented in the grey box. **(C)** Top candidates of the SAINTexpress analysis from 3 independent experiments for each condition. The average number of spectra (AvgSpec), the relative abundance and BFDR are represented. Among the top proteins, R2D2 is the second interactant of Dicer-2 for all three GFP::Dicer-2 lines and independent of the infection. **(D)** GO:biological processes enrichment analysis of GFP::Dicer-2^WT^ interactome proteins in mock-infected condition. **(E)** Global interaction network obtained with STRING/Cytoscape from the top 15% Dicer-2 partner proteins (i.e. top 44 proteins).

Dicer-2 forms a heterodimer with the dsRBP R2D2, and this association is essential for several steps in the siRNA biogenesis. Although Dicer-2 is able to cleave pre-miRNAs *in vitro*, R2D2 and inorganic phosphate restrict the specificity of Dicer-2, preventing pre-miRNA cleavage (Cenik *et al*, 2011; Hartig & Förstemann, 2011). R2D2 further amplifies this distinction between Dicer-1 and Dicer-2 by localizing Dicer-2 to cytoplasmic foci, called D2 bodies, where endo-siRNAs will be cleaved, away from the pre-miRNAs (Nishida *et al*, 2013). After siRNA processing by Dicer-2, R2D2 is then also required for exo-siRNA loading onto AGO2 (Liu *et al*, 2003, 2006; Marques *et al*, 2010), which is then stabilized by the Hsc70/Hsp90 chaperone machinery (Iwasaki *et al*, 2010; Miyoshi *et al*, 2010; Iwasaki *et al*, 2015). One strand of the siRNA duplex is then discarded, such that the remaining strand can guide the RISC complex to the complementary target RNA for silencing. For this step, R2D2 is needed again, as it functions as a protein sensor for thermodynamic differences in the base-pairing stabilities of the 5’end of the siRNAs. Thus, it allows the siRNA loading onto AGO2 in a specific orientation, thereby determining which strand will be discarded and which one will serve as the guide (Tomari *et al*, 2004; Yamaguchi *et al*, 2022).

Another dsRBP, the TRBP drosophila homologue Loquacious (Loqs), plays an important role in the determination of Dicer-2 specificity. Because of alternative splicing, there are four distinct Loqs isoforms, with specific activities in the Dicer-1-dependent miRNA biogenesis pathway or the Dicer-2-dependent endo-siRNA pathway. While the function of Loqs-PC remains unknown, Loqs-PA and Loqs-PB interact with Dicer-1 for miRNA biogenesis, and Loqs-PD interacts with Dicer-2 for the biogenesis of endo-siRNAs (Förstemann *et al*, 2005; Jiang *et al*, 2005; Saito *et al*, 2005; Hartig *et al*, 2009; Zhou *et al*, 2009; Marques *et al*, 2010). Intriguingly however, Loqs-PD is not required for the targeting of viral dsRNA for the viruses tested (Marques *et al*, 2013). This last isoform is able to modulate the termini dependence of Dicer-2 by enabling the cleavage of sub-optimal substrates such as dsRNA with blocked, structured, or frayed ends (Sinha *et al*, 2015). This modulation is not achieved by changing the cleavage mode (i.e. processive and distributive), but rather by affecting the probability for Dicer-2 to cleave the sub-optimal substrate (Naganuma *et al*, 2021).

In order to get a clearer picture of the protein network associated with Dicer-2 during viral infection and test the impact of Dicer-2 partners in antiviral immunity, we used an interactomics approach through immunoprecipitation followed by mass spectrometry (IP-MS), in a fly line expressing the wild-type version of Dicer-2 fused to a GFP tag, GFP::Dicer-2^WT^. As we were mostly interested in the early steps of the RNAi pathway (i.e. the viral dsRNA sensing by Dicer-2), we also used two Dicer-2 mutants that were able to sense the dsRNA but unable to process it. One of those mutants expresses a GFP-tagged version of Dicer-2 with the G31R mutation on the Hel1 domain, called GFP::Dicer-2^Hel1^. This Dicer-2 mutant is able to process a 3’overhang dsRNA substrate using distributive dicing, but not a blunt dsRNA substrate, which requires ATP hydrolysis for processive dicing. It has been described in previously published work, together with the impact of this mutation on endo- and exo-siRNA production (Donelick *et al*, 2020). We also used another Dicer-2 mutant fly line, GFP::Dicer-2^RNaseIII^, with two mutations in the RNase III domains, in the hope of identifying more transient interactions, as this mutant is able to bind dsRNA but not cleave it. Using these tools, we report the identification and functional characterization of novel interactants for Dicer-2. Amongst them, we observed proteins such as Me31B and eIF4E1, for which the interaction with Dicer-2 is RNA-independent, whereas the protein Syncrip (Syp) interacts with Dicer-2 in an RNA-dependent manner. By performing two RNAi screens, both in S2 cell culture and *in vivo*, we have highlighted several of those proteins as having an impact on viral DCV infection. In particular, the protein Rasputin (Rin) has an antiviral impact on DCV RNA load both *ex vivo* and *in vivo*, as well as an impact on the survival after DCV infection *in vivo*.

## RESULTS

### Definition of the Dicer-2 interactome *in vivo*

To study the dynamics of the protein network surrounding Dicer-2 *in vivo* in response to viral infection, we complemented *dicer-2* null mutant fly lines with WT or mutant versions of GFP::Dicer-2 and injected them with either TRIS (mock-infection) or DCV. After immunoprecipitation of the different GFP::Dicer-2 versions, their protein partners were then identified by LC-MS/MS (**Figure 1A & B, Supplementary Figure S1A & B**). The Hel1 and RNase III mutants should provide an overview of the interactome of Dicer-2 at the early steps of dsRNA recognition. All experiments were performed in adult flies, and the ability of the wild-type version of Dicer-2 (GFP::Dicer-2^WT^) to rescue the *dicer-2* null mutation was demonstrated in previous studies (Kemp *et al*, 2013; Girardi *et al*, 2015; Donelick *et al*, 2020). Furthermore, two control fly lines were used to determine non-specific interactants, both expressing endogenous *dicer-2* normally: a wild-type CantonS line (Ctrl-WT) and a transgenic line expressing GFP ubiquitously (Ctrl-GFP). Of note, the level of expression of GFP::Dicer-2 in the complemented lines is comparable to the endogenous expression of Dicer-2 in control lines (**Supplementary Figure S1A**).

In total, 2511 protein hits were identified by LC-MS/MS across all samples. As our approach uses several versions of Dicer-2, a first analysis was performed using the SAINTexpress tool (Teo *et al*, 2014) to have a global overview of the Dicer-2 partners. This allowed us to create a list of 288 proteins that were significantly enriched compared to the controls overall (**Supplementary Figure S1C and Supplementary Table 1**). Visualization of this analysis using Prohits-viz (Knight *et al*, 2017) showed that the bait protein Dicer-2 is enriched in all GFP::Dicer-2 lines as expected (**Figure 1C**). Of note, specific peptides of the different GFP::Dicer-2 fusions were found in the MS data, confirming the presence of Dicer-2^WT^, Dicer-2^Hel1^ and Dicer-2^RNaseIII^ in the complemented lines (data not shown). Moreover, we can observe amongst the main interactants several proteins known to be involved in RNAi, like R2D2, a known cofactor of Dicer-2, AGO2, Loqs and CRIF (**Supplementary Table 1**). These results, consistent with the literature, validate the reliability of the approach (Liu *et al*, 2003; Czech *et al*, 2008; Miyoshi *et al*, 2010; Cernilogar *et al*, 2011; Hartig & Förstemann, 2011; Lim *et al*, 2014).

### Global analysis of the protein network surrounding Dicer-2 *in vivo*

Analysis of the biological processes associated with the GFP::Dicer-2^WT^ interactome revealed five statistically enriched GO terms (**Figure 1D, Supplementary Table 1**). As expected, this Dicer-2 interactome reveals an enrichment in proteins linked to small RNA pathways and antiviral defense. After ranking candidates obtained with SAINTexpress based on Bayesian false discovery rate (BDFR) values and fold-change between all the GFP::Dicer-2 lines, we selected the top 15% proteins (i.e. the 44 top candidates) to establish a global network using the STRING database (**Figure 1E**). Amongst them, four proteins involved in RNAi were identified. Moreover, we observed a node of proteins involved in the response to external stimulus that interact with each other. One of those proteins, NorpA, is a phospholipase C enzyme which genetically interacts with the NF-κB pathway regulating neuronal cell death in drosophila (Chinchore *et al*, 2012). Of note, several members of this node have been reported to interact physically (Ye *et al*, 2018; Chen & Montell, 2020; Chen *et al*, 2021). The GO term “Ribonucleoprotein complex” was also highlighted both in the GO term enrichment analysis and in the STRING analysis. Finally, we found several proteins involved in the spliceosome complex (e.g. Syp, Lost, Rump, Exu). These results may point to hitherto uncharacterized roles of Dicer-2.

### Dicer-2 point mutations reveal specific interaction profiles in mock-infected and DCV-infected samples

We then focused our analysis on the respective impacts of the Dicer-2 helicase and RNaseIII domain mutations and of the DCV infection on this interactome, as studying how it is modulated in these different conditions could help us understand better the roles of the different Dicer-2 interactants identified above. To this aim and in addition to the global analysis, we performed separate statistical analyses to compare each GFP::Dicer-2 line to each other. To achieve this, we used a negative-binomial test to identify proteins enriched in each GFP::Dicer-2 line compared to the control lines (Ctrl-CS and Crtl-GFP), with a fold-change>2 and an adjusted *p*-value<0.01. Using the same approach, we then determined the impact of DCV infection on the interactome of Dicer-2. This allowed us to group the different proteins interacting with Dicer-2 into categories depending on their enrichment in the different GFP::Dicer-2 lines and in DCV-infected samples (**Figure 2A-C, Supplementary Figure S2A-D**). We can observe that 146 proteins appear to be enriched in all GFP-Dicer-2 lines, including 25 proteins that were also enriched in the DCV-infected samples.

**Figure 2:**
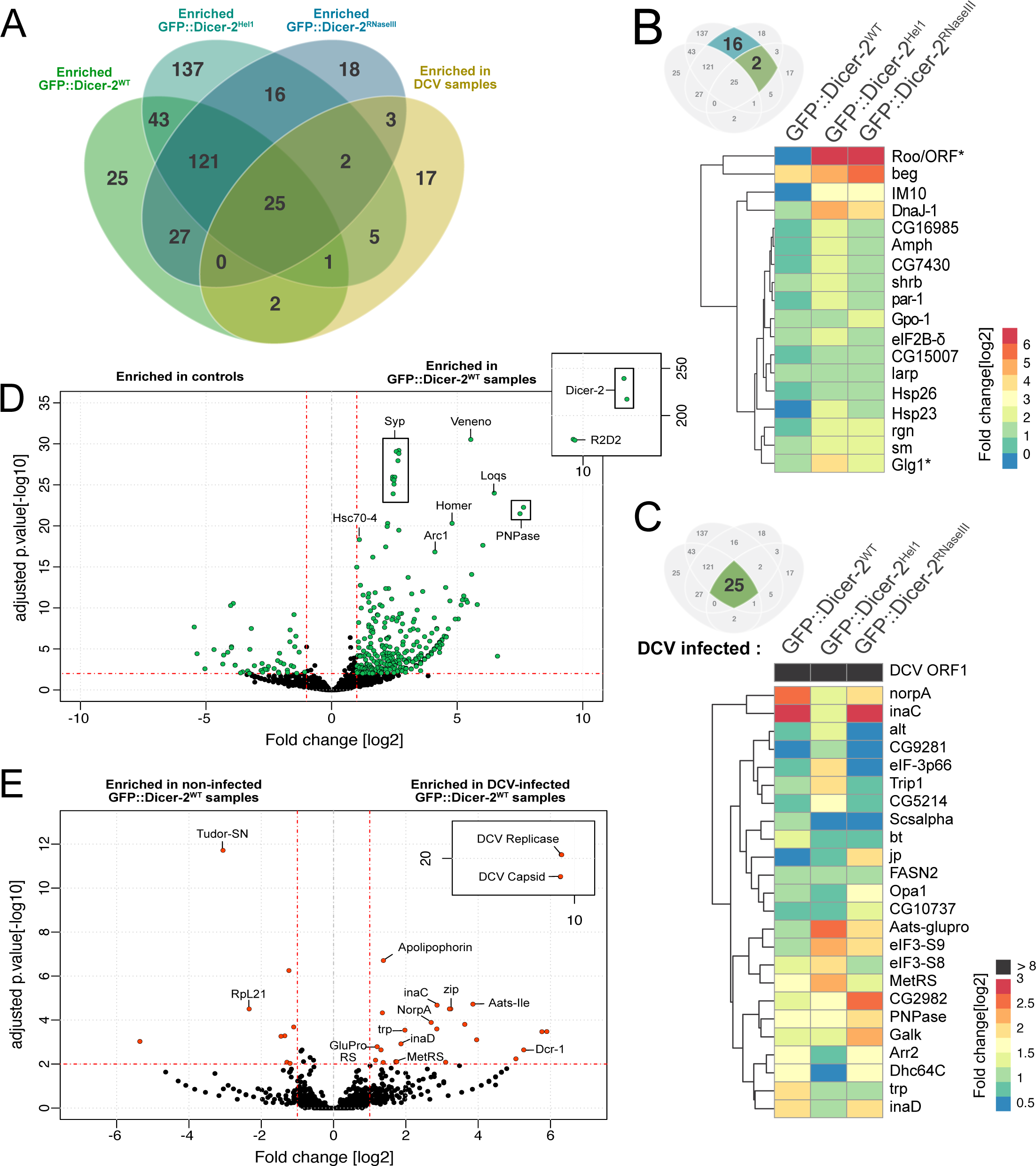
Impact of the Dicer-2 mutations on the RNP network during DCV infection. **(A)** Venn diagram showing the number of proteins identified in each GFP::Dicer-2 line in mock-infected and DCV-infected adult flies. Candidates are selected with a fold-change > 2 and an adjusted *p-*value < 0.01. **(B)** Heatmap representing the fold-change in comparison to the controls of the 18 proteins enriched only in the two GFP::Dicer-2 mutants (Hel1 and RNAseIII). 16 proteins are stable in mock-infected and DCV-infected samples. Two proteins, Roo/ORF and Glg1, are specifically enriched during DCV-infection and indicated with an asterisk. Log2 fold-change for each protein and each genotype is represented. **(C)** Heatmap representing the fold-change in comparison to the controls of the 25 proteins enriched with all GFP::Dicer-2 lines during DCV infection. The set of proteins is composed of 24 Drosophila proteins and DCV RdRp (ORF1). Log2 fold-change for each protein and each genotype during DCV infection is represented. **(D)** Volcano plot representing the fold changes and adjusted *p*-value of the Dicer-2 partners in GFP::Dicer-2^WT^ lines in mock- and DCV-infected conditions *versus* the control lines (Crtl-CS and Crtl-GFP). All the proteins with fold change > 2 and an adjusted *p*-value < 0.01 are represented in green. **(E)** Volcano plot representing the fold changes and adjusted *p*-value of the GFP::Dicer-2^WT^ partners in mock-infected *versus* DCV-infected adult flies. All the proteins with fold change > 2 and an adjusted *p*-value < 0.01 are represented in orange.

For every GFP::Dicer-2 line, R2D2 was always one of the top partners of Dicer-2, consistent with the presence of a stable heterodimer Dicer-2/R2D2 *in vivo* (Liu *et al*, 2003). This is illustrated in the volcano plots comparing each GFP::Dicer-2 line *versus* the controls (**Figure 2D, example of GFP::Dicer-2^WT^**). Among the most enriched proteins in GFP::Dicer-2^WT^ samples, the Tudor protein Veneno (CG9864) and the chaperonin Hsc70-4 are involved in small RNA pathways (Iwasaki *et al*, 2010; Handler *et al*, 2011; Joosten *et al*, 2019). We can also observe the presence of Syncrip (Syp), and another candidate from the global analysis, Polynucleotide phosphorylase (PNPase). PNPase is involved in RNA import into mitochondria and RNAs degradation during apoptosis in mammals (Portnoy *et al*, 2008; Liu *et al*, 2018a). Finally, Arc1 is a Gag-like retrotransposon able, in mammals, to self-assemble into capsids to encapsulate RNA and release it in extracellular vesicles (Pastuzyn *et al*, 2018). All these proteins represent the Dicer-2^WT^ network *in vivo*. Interestingly, GFP::Dicer-2^Hel1^ interacts specifically with 142 proteins, including 137 in mock-conditions whereas GFP::Dicer-2^WT^ and GFP::Dicer-2^RNaseIII^ have only 27 and 21 specific interactants respectively (**Figure 2A**). As mentioned above, the GFP::Dicer-2^Hel1^ mutant is unable to perform the more efficient processive dicing and can only perform distributive dicing, we therefore expect it to interact longer with the viral RNA, as this is a slower process. This result thus suggests that some proteins could interact transiently with Dicer-2 or only during distributive dicing, and be stabilized with Dicer-2^Hel1^.

To better characterize the early steps of Dicer-2 sensing, we focused on the 18 proteins interacting specifically with the two mutant lines GFP::Dicer-2^Hel1^ and GFP::Dicer-2^RNaseIII^, including two proteins, Roo/ORF and Glg1, that were also enriched in DCV infected conditions. These proteins are represented on a heatmap on **Figure 2B**, and studying them could help us understand the ribonucleoprotein complexes involved in the molecular mechanism of dicing (**Supplementary Figure S2B**). Taken together, these results illustrate the differences in the protein networks of the two Dicer-2 mutants compared to GFP::Dicer-2^WT^. Moreover, the legitimacy of the top 15% candidates chosen after the global analysis is further confirmed by the fact that out of those 44 proteins, 36 of them are also found in the 146 proteins enriched in all GFP::Dicer-2 samples (**Supplementary Figure S2C**). Out of the 8 proteins remaining, 4 were enriched in two of the different GFP::Dicer-2 lines. Most candidates on this list have therefore been highlighted in four different analyses, using two different methods.

Next, we analyzed the impact of the DCV infection on Dicer-2 networks by studying the proteins enriched in the DCV-infected samples compared to the mock-infected samples (**Figure 2E**, example of DCV-infected GFP::Dicer-2^WT^ samples compared to mock-infected GFP::Dicer-2^WT^ samples). By adding this data to the one obtained during the previous analysis, we were able to categorize the different proteins depending on their enrichment in the different GFP::Dicer-2 lines and/or the DCV-infected samples (**Figure 2C, Supplementary Figure S2D**). Amongst those 25 proteins, 11 are present in the top 15% list of candidates from the global analysis. The interactions between Dicer-2 and those proteins seem to increase after infection with DCV, which could indicate a connection of those proteins with the antiviral RNAi function of Dicer-2 (**Figure 2C**). Of note, these interactants include the viral RNA-dependent RNA polymerase (RdRp) from DCV (ORF1). This is most likely due to Dicer-2 and the DCV RdRp being present on the same viral RNA molecule at the same time. GO term analysis for “Biological processes” terms shows that the GFP::Dicer-2 lines are enriched for proteins involved in small RNA pathways. These GO terms are highly enriched in the GFP::Dicer-2^Hel1^ suggesting a stabilization of the antiviral Dicer-2 interactome in this genetic background (**Supplementary Figure S2E**).

### Validation of the interaction between Dicer-2 and new protein partners

In order to validate the accuracy of the above-mentioned analyses, we decided to focus on a set of candidates, namely the candidates of the global analysis, and confirm that they do indeed interact with Dicer-2. We therefore studied the interactions between these main candidates and GFP::Dicer-2 by anti-GFP immunoprecipitation and western blot (**Figure 3**). This allowed us to confirm the interaction of Me31B, Rump, eIF4E1 and Syp with GFP::Dicer-2^WT^ as well as the two mutants (**Figure 3A**). These proteins are known to be part of the same RNA-protein complexes (Nakamura *et al*, 2004; Igreja & Izaurralde, 2011; McDermott *et al*, 2012; Wang *et al*, 2017). The interaction between Dicer-2 and its partners does not appear to be dependent on DCV infection (**Figure 3B**). On the contrary, after infection with DCV we can observe a weaker co-immunoprecipitation of the different candidates, which could suggest that Dicer-2 associates with mRNP complexes linked to other functions of Dicer-2, like translation repression. Overall, these experiments confirm the results obtained with the global MS analysis. We note that the interactions seem stronger with the GFP::Dicer-2^Hel1^ mutant, which is also consistent with the MS analysis, in which more spectral counts are observed with the GFP::Dicer-2^Hel1^ line (**Figure 3A, Supplementary Figure S1C**). This could be due to a stabilization of these interactions in the GFP::Dicer-2^Hel1^ mutant. Moreover, in the raw MS data very few spectra for Syp, Rump and eIF4E1 were identified with the Ctrl-GFP and Ctrl-WT lines, while more spectra were identified in the controls for Me31B (**Supplementary Table 1**). The fact that Me31B was, although in lower amounts than for the GFP::Dicer-2 lines, also found in the Ctrl-GFP IP (**Figure 3B**) is therefore consistent with the MS results. Moreover, these interactions do not seem to require DCV infection to occur, which we also observed in the MS data, as spectra for those proteins were also identified in the non-infected samples (**Supplementary Figure S1C, Supplementary Table 1**).

**Figure 3:**
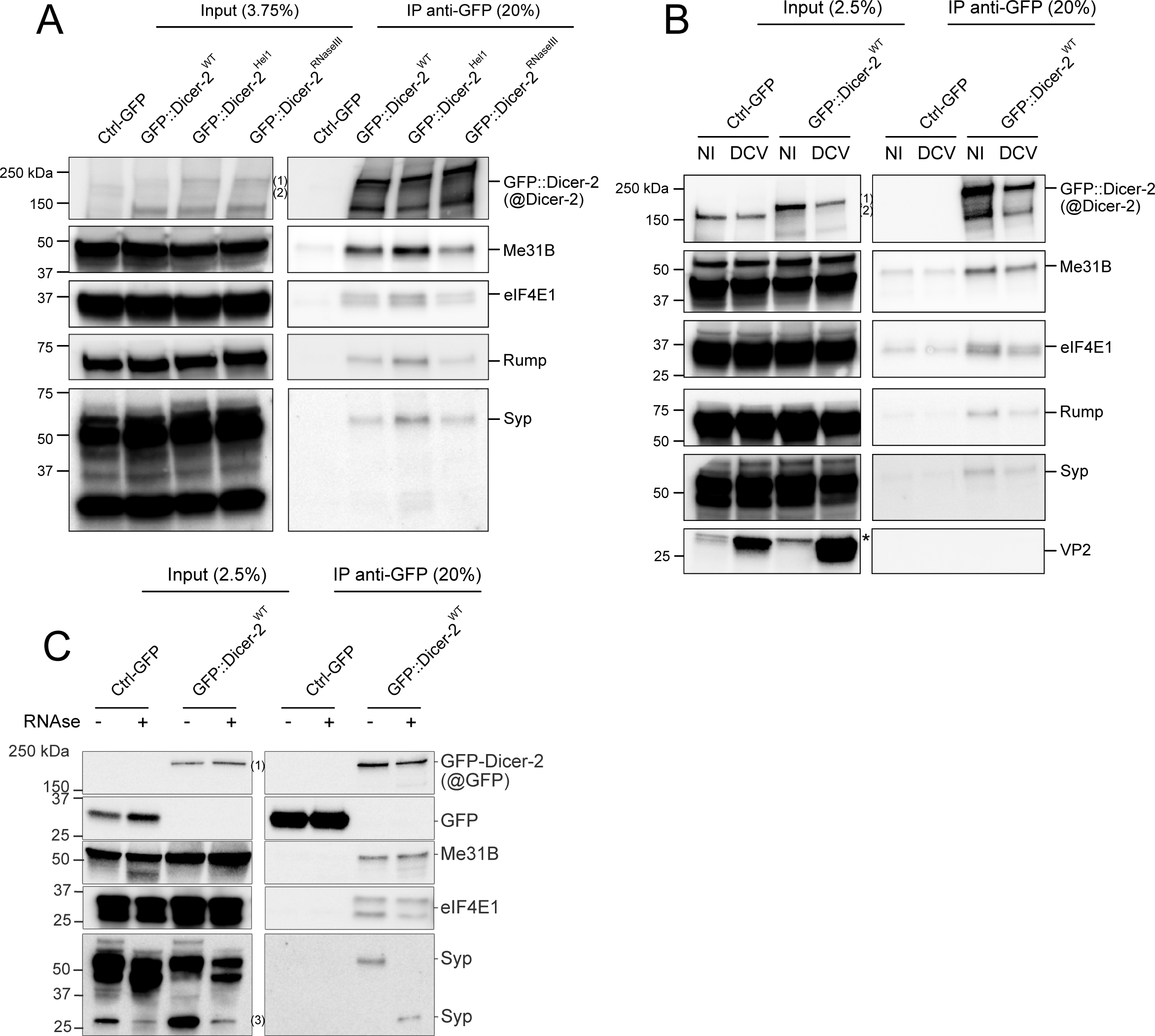
Validation of the Dicer-2 interactome. Results of the mass spectrometry analysis were confirmed by immunoblot after immunoprecipitation with anti-GFP beads. (1) Size of GFP::Dicer-2; (2) Size of endogenous Dicer-2; (3): Lower isoform of Syp. The asterisk (*) represent a band non-specific to the anti-VP2 antibody. The experiment was performed using: **(A)** GFP::Dicer-2^WT^, GFP::Dicer-2^Hel1^ and GFP::Dicer-2^RNAseIII^ in comparison to the Ctrl-GFP line; **(B)** non-infected (NI) or DCV-infected GFP::Dicer-2^WT^ flies in comparison to the Ctrl-GFP line; **(C)** GFP::Dicer-2^WT^ in comparison to the Ctrl-GFP line, after treatment (+) or not (-) with RNase A.

### RNA participates in the interaction between Dicer-2 and some of its associated proteins

As we decided to perform the immunoprecipitations in low stringency conditions, the IP-MS analysis allowed us to identify a high number of interactions with Dicer-2. Those could be direct protein-protein interactions, but also indirect interactions through other proteins or RNAs, as we were interested in studying the complexes as a whole. To check if the interaction of Dicer-2 with its partners was RNA-dependent, we performed IPs of GFP::Dicer-2 with and without RNase A treatment (Figure 3C, Supplementary Figures S3A & S3B). We observed that Dicer-2 interacts with Me31B and eIF4E1 in an RNA-independent manner, as both partners co-immunoprecipitate with Dicer-2 with and without RNase A (**Figure 3C**). By contrast, the interaction between Rump and GFP::Dicer-2 also seems to be RNA-dependent, as we can observe a weaker co-immunoprecipitation of Rump after RNase treatment (**Supplementary Figure S3B**). Most strikingly, we observed a modification of the interaction of GFP::Dicer-2 with the protein Syncrip (Syp), which has several isoforms (McDermott *et al*, 2012). Indeed, the high molecular weight isoform of Syp interacting with GFP::Dicer-2 was displaced upon RNase treatment for a shorter isoform of Syp (**Figure 3C, Supplementary Figure S3B**). Of note, the RNase A treatment also impacts Syp isoforms in the total protein extract.

To expand on these results, we performed a second IP-MS experiment on GFP::Dicer-2^WT^ with or without RNase A treatment, wherein adult flies expressing either GFP alone (Ctrl-GFP) or GFP::Dicer-2 ^WT^ were injected with TRIS (mock-infection) or DCV (**Supplementary Figure S3C**). Four independent statistical analyses were performed using the same strategy as described above, in order to compare each condition (mock-infected or DCV-infected and/or treated with RNase A) to its corresponding control. In total, this new set of data allowed us to highlight 448 proteins interacting with GFP::Dicer-2^WT^ in the different conditions compared to the Ctrl-GFP flies. Among them, we were able to identify most of the candidates from the previous MS experiment analysis in the non-RNase treated samples, which demonstrates the reproducibility of the two IP-MS experiments. As expected, Rump and Syp, which were shown to be RNA-dependent (**Figure 3C, Supplementary Figure S3B**), were not enriched in any of the RNase-treated conditions (**Supplementary Figure S3C**), while Dicer-2 and its known cofactor R2D2 were found in all four conditions. Taken together, these results allow us to characterize the RNA-dependence or independence of the interactions between the top candidates and Dicer-2. Globally, we found that 189 out of the 317 proteins (60%) highlighted by these analyses were identified in at least one RNAse-treated condition (**Supplementary Figure S3C**). These results correlate with the results obtained by Nadimpalli *et al*. (Nadimpalli *et al*, 2022), which also found that a high number (81%) of Dicer-2 interactants were RNA-independent in the early drosophila embryo.

### Identification of new proviral and antiviral factors in drosophila S2 cells

Overall, we established a global interactome of Dicer-2 *in vivo*, which may include previously unknown components of the antiviral RNAi pathway or regulators of viral infection. Therefore, we next assessed the impact of these interactants on viral infection by DCV. The top 15% candidates of the global SAINT analysis were therefore subjected to an RNAi screen in S2 cells, to test their function during viral infection. To this end, the expression of the different candidate genes was inhibited in S2 cells using dsRNAs from DRSC Harvard prior to infection by DCV and monitoring of the viral RNA load 20h later (**Figure 4A**). Whenever possible, two different dsRNAs were used for each candidate gene in order to discount off-target effects, and the dsRNAs targeting RACK1 and AGO2 were used as a proviral and antiviral controls respectively. The data were normalized and analyzed using a linear mixed effect model, and the viral RNA loads of the candidates were compared to that of the dsLacZ control (**Figure 4B**). Of note, knock-down of AGO2 does not exhibit a significative antiviral response, probably due to a conflict with the RNA interference pathway that requires the AGO2 protein. However, we found that some of the candidates, eIF4G1 (2/2 dsRNAs) and Rin (1/2 dsRNA), exhibited a significant antiviral phenotype against DCV infection. Moreover, knock down of *lig* led to a significantly decreased DCV RNA load suggesting a proviral function of the Lig protein (1/2 dsRNA, although the viral load of both dsRNAs are very similar). Moreover, we observed that *rump* knock down (KD) led to one of the highest viral load amongst the candidates tested, although it did not reach statistical significance in our assay conditions (**Figure 4B**). Rump is a hnRNP M homologue that binds to *nos* and *osk* mRNAs and has hitherto not been implicated in antiviral immunity. The other proteins for which the interaction with Dicer-2 was validated (Me31B, eIF4E1 and Syp) do not seem to have any impact on DCV RNA load at all, suggesting that they might be involved in the other functions of Dicer-2. Overall, a high proportion of the candidate genes show an increase of DCV viral load after KD although not statistically significant, suggesting that they could still play a minor role in antiviral immunity in drosophila S2 cells.

**Figure 4:**
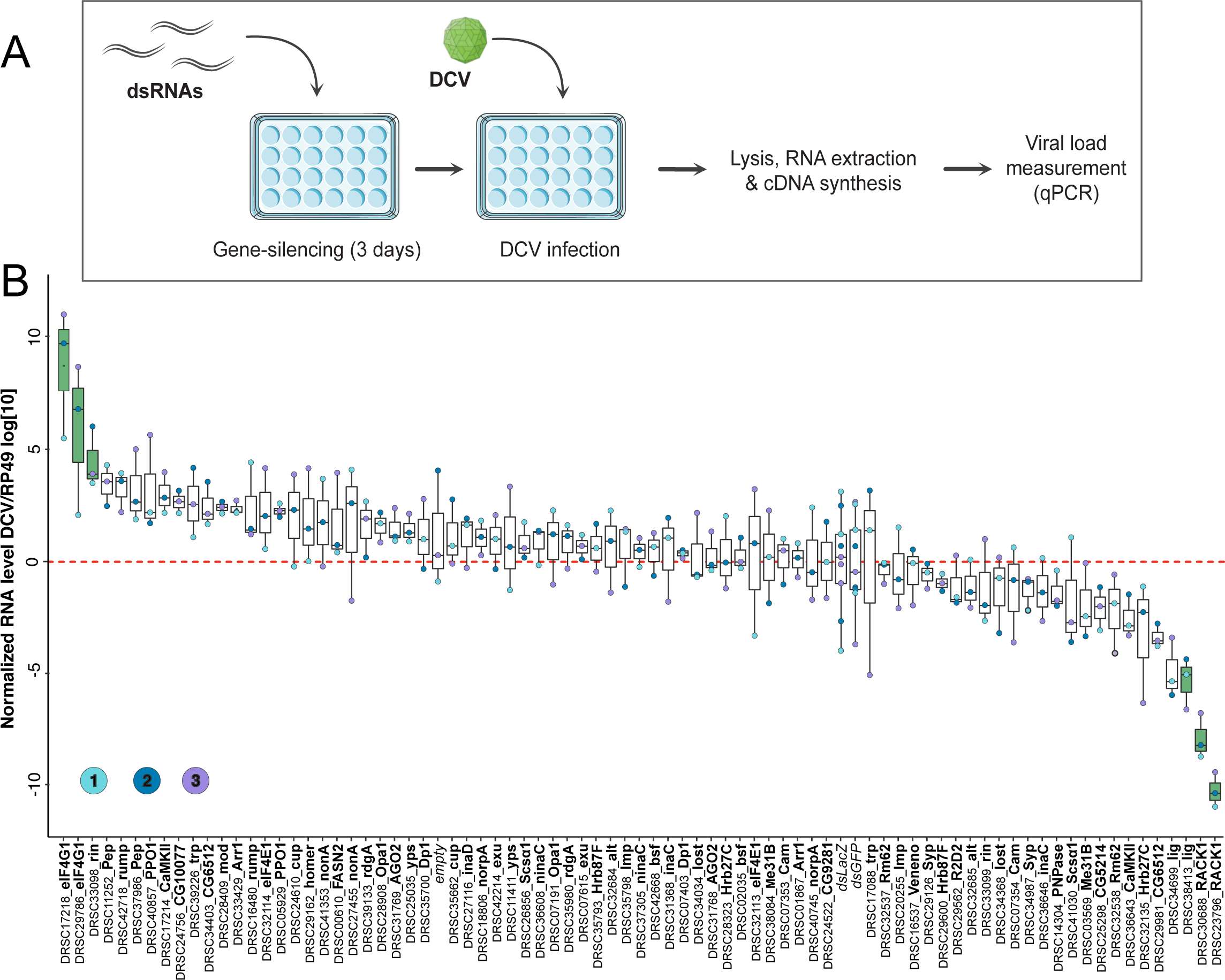
Role of the candidates in antiviral immunity in drosophila S2 cells. **(A)** Schematic representation of the experiment procedure. Knock-downs (KD) of gene expression were performed by soaking drosophila S2 cells with dsRNA targeting the candidate genes during 3 days. On day 3, cells were infected with DCV at 0.001 MOI for 20h. **(B)** Box plot representing the DCV viral RNA load of S2 cells after KD of the top 15% candidates followed by DCV infection. The dsRNAs targeting candidate genes are represented with the DRSC number and the gene name. Negative controls are infected S2 cells that were not treated with any dsRNA (empty), dsRNA against GFP, dsRNA against LacZ. In adition, dsRACK1 was used as a proviral control. The DCV RNA level is normalised to the housekeeping mRNA *RP49*. The three independent experiments are indicated by the color of the dots. The significant antiviral and proviral candidates are in green with a *p-*value < 0.05.

### Identification of new proviral and antiviral factors *in vivo*

We next evaluated the impact of a subset of the interactants, corresponding to the top 10% of the global analysis but excluding candidates for which the KD was lethal *in vivo* (e.g. eIF4G1), on survival after DCV infection in the context of the whole organisms. A knock-down (KD) of the different candidate genes was induced through temperature shift in adult flies using the [*actin*-Gal4; *tub*-Gal80^TS^] system, and the flies were then injected with DCV (**Figure 5A**).

**Figure 5:**
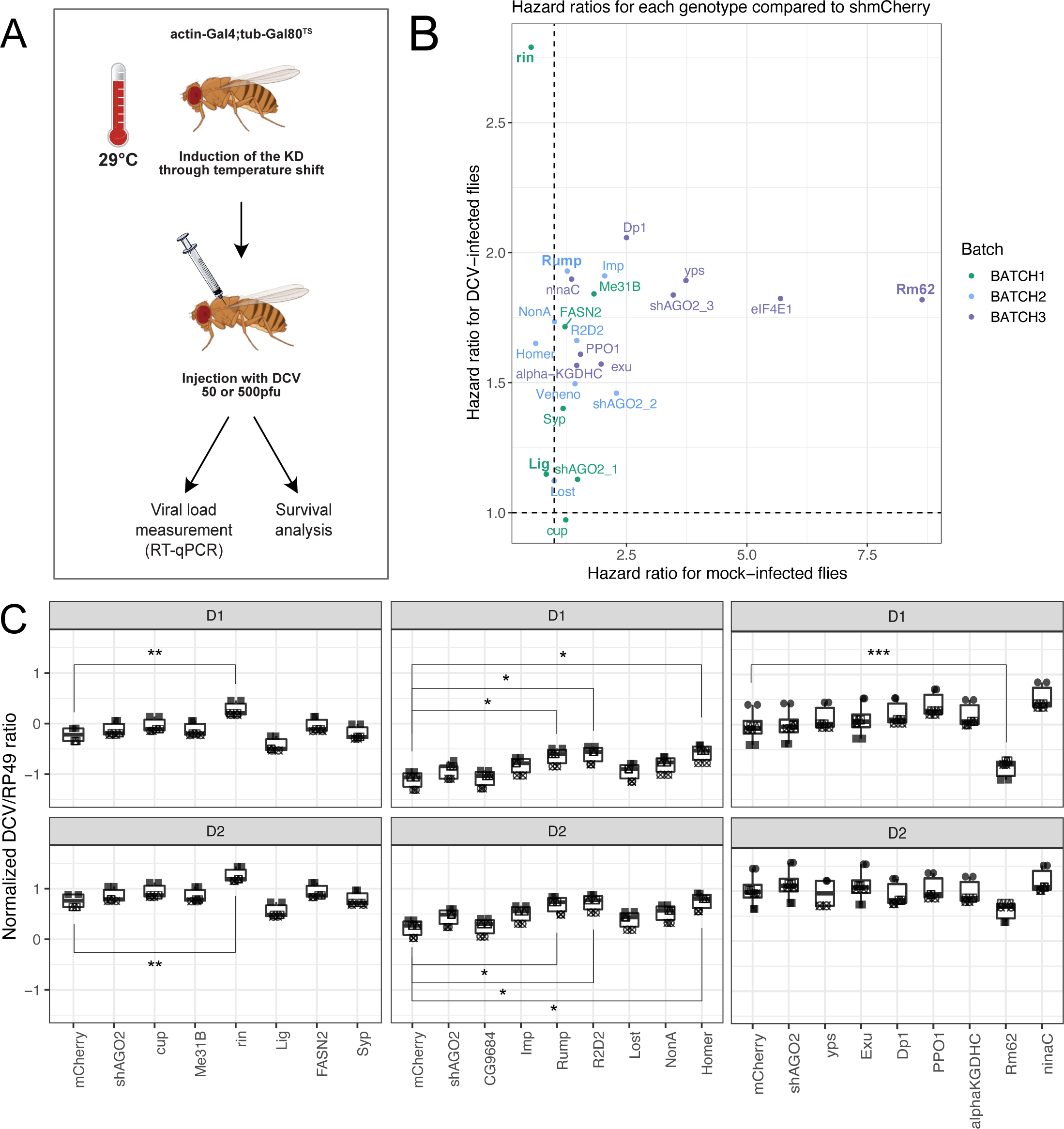
Role of the candidates in antiviral immunity in adult drosophila flies. **(A)** Schematic representation of the experiment procedure. Knock-down (KD) of gene expression were established by crossing inducible UAS-dsRNA for each candidate gene with an ubiquitous Gal4 driver (actin5C-Gal4) under the control of thermosensitive ubiquitous Gal80 (tub-Gal80^TS^) during five days at 29°C. Adult flies were injected with DCV 50pfu and analysed for survival or DCV 500pfu followed by RT-qPCR at days 1 and 2 post-infection. **(B)** Hazard ratios of the different UAS-dsRNA lines after mock-infection or DCV-infection. **(C)** Box plot representing the DCV viral RNA load after infection of DCV 500 pfu at day 1 and day 2 post-infection. Triplicate experiments represented in three batches. Significant viral load compared to shmCherry line are represented.

Survival was assessed by counting flies every day for 20 days and computing the hazard ratios for each KD fly line, which corresponds to the chances of the different KD flies dying compared to the control shmCherry flies (**Figure 5B, Supplementary Figure S4**). In addition, viral RNA load was also measured by RT-qPCR at 1 and 2 dpi after injection (**Figure 5C**).

Unfortunately, the strongest candidate highlighted by the *ex vivo* screen, eIF4G1, could not be tested *in vivo* as its knockdown is lethal in the whole fly. The most striking impact on viral infection *in vivo* was observed for Rin. Indeed, although the hazard ratio for Rin in mock-infected flies is close to 1, meaning that the injection of TRIS alone does not impact survival in Rin KD flies, the hazard ratio for DCV-infected Rin KD flies is significantly higher than 1, meaning that Rin KD has a negative impact on survival after DCV infection (**Figure 5B**). This correlates with the higher DCV RNA load observed *in vivo* (**Figure 5C**), which was already observed *ex vivo* (**Figure 4B**). Taken together, these data strongly suggest that Rin is involved in the antiviral response against DCV in *Drosophila melanogaster*.

Similarly, the hazard ratio for Rump KD flies is higher than 1 after infection with DCV but not TRIS (**Figure 5B**), and is correlated with a higher DCV RNA load *in vivo* (**Figure 5C**), confirming the trend observed *ex vivo* (**Figure 4B**). Thus, Rump may also be involved in the antiviral reponse against DCV. We also tested the role of Lig, however *in vivo* data monitoring survival (**Figure 5B**) or viral RNA load (**Figure 5C**), did not confirm the proviral effect observed in cell culture (**Figure 4B**). Amongst the other candidates tested, we observed a strong proviral impact of another candidate, Rm62, on DCV RNA load (**Figure 5C**). However, Rm62 KD also highly impacts survival in non-infected conditions, which suggests that Rm62 impacts the general stress response after injection.

## DISCUSSION

The crucial role of a few known protein partners of Dicer-2 in regulating or facilitating its activity has been established (Cenik *et al*, 2011; Hartig & Förstemann, 2011; Hansen *et al*, 2019). Yet, we are far from truly understanding the many functions of this multifunctional protein. In order to establish the protein network of Dicer-2 *in vivo* in different conditions, we have used different tagged GFP::Dicer-2 constructs and an IP-MS approach. We have confirmed the reliability of this approach by the identification of the usual key players of RNAi, also found in other interactomes of RNAi proteins (Frohn *et al*, 2012; Gerbasi *et al*, 2010; Joosten *et al*, 2021; Montavon *et al*, 2021; Varjak *et al*, 2020), e.g. R2D2, Ago1, AGO2 and Loqs. Moreover, we were also able to observe recurring interactions compared to the networks established in other studies. Indeed, we were able to confirm the interaction of Me31B, also known as DDX6, with Dicer-2 in drosophila *in vivo*, and this protein was also identified in mammalian cells as a potential interactant of AGO2 (Frohn *et al*, 2012). The protein Larp4B, which was found in the list of 146 proteins interacting with all three GFP::Dicer-2 lines used, provides another example. This nucleic acid binding protein interacts with Dicer-2 in the embryo and has been shown to be antiviral against DCV, CrPV, SINV and VSV (Nadimpalli *et al*, 2022; Pennemann *et al*, 2021). Moreover, we have been able to identify proteins that were shown to interact with Dicer *ex vivo* in other species, e.g. DHX9/Mle and proteins from the Heat shock protein family 70 (Montavon *et al*, 2021; Varjak *et al*, 2020).

By contrast with the previous interactomes of Dicer-2 in other species this is, to our knowledge, the first interactome of Dicer-2 performed *in vivo* in adult flies. Moreover, in addition to a global analysis of the protein network of Dicer-2, we have also studied the impact of two point mutations affecting key domains of the protein. This has allowed us to highlight proteins interacting with only one, two, or all GFP::Dicer-2 samples. Our results reveal a list of proteins interacting with all three GFP::Dicer-2 lines, which indicates a very stable interaction, not disturbed by those specific mutations, and reproducible as it was observed with three different GFP::Dicer-2 fly lines. Therefore, by choosing to study the list of 44 top candidates from the global analysis, we have biased our study towards very stable interactions with Dicer-2. In addition, we also identified proteins interacting only with the WT version of GFP::Dicer-2. These proteins may correspond to factors of a functional Dicer-2 complex, actively producing siRNAs. On the contrary, we identified proteins interacting only with the two mutants, which sequestrate the dsRNA in an unproductive complex (Donelick *et al*, 2020). These proteins may correspond to factors interacting with Dicer-2 in a transient manner. In particular, the 142 proteins found only in the GFP::Dicer-2^Hel1^ samples (**Figure 2A, Supplementary Figure S2A**) could be a consequence of the impaired dsRNA processing of this mutant, allowing the stabilization of these interactions. For example, we have noticed a stronger interaction of the two GFP::Dicer-2 mutants with a transposable element called Roo/ORF, which is not present for the WT version of Dicer-2. Keeping in mind that transposable elements participate in the reverse-transcription of viral RNA and have been proposed to play a role in systemic antiviral RNAi (Goic *et al*, 2013), the study of Roo/ORF could prove interesting. We provide a list of proteins whose interaction with Dicer-2 depends on the presence of a functional helicase or RNase III domain. Finally, one of the newly identified antiviral candidates from the global analysis, Rin, with an *adjp*-value<0.01, was not significantly enriched in the GFP::Dicer-2^RNaseIII^ mutant in the fly line-specific analysis. Therefore, the RNase III domain may be necessary for this interaction, although this needs to be confirmed biochemically.

In the global analysis of the Dicer-2 network, although obtained from a mixed population of adult male and female flies, many proteins are well described for their functions during drosophila oogenesis, including three of the proteins for which the interaction with Dicer-2 was confirmed by WB: Me31B, eIF4E1 and Rump. By looking deeper into those candidates, we noticed that most of them (e.g. Dp1, Lost, Cup, eIF4E1, Me31B, Imp, Syp, Bsf, Rump, Exu & Yps) were associated with mRNA localization regulation mechanisms, like the regulation of *nanos* or *oskar* mRNA translation. Indeed, Me31B has been shown to form a complex with Tral, eIF4E1, Cup and PABP involved in RNP-mediated translational repression of maternal mRNAs during oogenesis and embryogenesis (Wang *et al*, 2017). It has also been shown to form a complex with Exu and Yps involved in the translational silencing of mRNAs (such as *oskar*) during their transport to the oocyte (Nakamura *et al*, 2001). The presence of eIF4E1, Cup, Exu and Yps in the top candidates of the global analysis could suggests that Dicer-2 could interact with those two complexes formed by Me31B. An impact of Dicer-2 in oogenesis could explain the decreased fertility of *dicer-2 null* mutants. Recently, Dicer-2 has been shown to form a complex with Ataxin-2/Tyf forming a noncanonical cytoplasmic polyadenylation complex (Nadimpalli *et al*, 2022). In addition, Rump and Lost, which interact together, have also been shown to be involved in mRNA regulation in the oocyte (Jain & Gavis, 2008; Sinsimer *et al*, 2011) and this is also the case for Syp (McDermott *et al*, 2012). This would not be the first time that an RNAi-independent role in mRNA regulation of Dicer-2 is reported, as previous studies have already reported an RNAi-independent role of Dicer-2 in the regulation of mRNA expression, and more specifically in the activation of the expression of *Toll* and *R2D2* mRNAs through cytoplasmic polyadenylation (Coll *et al*, 2018; Wang *et al*, 2015). Interestingly, another small RNA factor, Aubergine (Aub), has been implicated in both mRNA localization regulation and Wispy-related mRNA regulation mechanisms (Dufourt *et al*, 2017) and has been implicated as a *nanos* mRNA localization factor in a mechanism implicating Rump (Becalska *et al*, 2011). If Dicer-2 does have a link with the regulation of mRNA expression during oogenesis, it would underline the interconnection between small RNA factors and RNAi-independent mRNA regulation mechanisms.

An important objective of this study was to analyze the impact of DCV infection on the protein network of Dicer-2, and the contribution of the interactants identified to the control of the infection. These experiments have highlighted four candidates (Rump, Lig, Rin and Rm62).

Rm62 is a member of the DDX5/Dbp2 subfamily of DEAD-box helicases (Xing *et al*, 2019, 5), which have been shown to be important in virus-host interactions (Rousseau & Meignin, 2020). In particular, some DEAD-box helicases play a role in RNAi and bind dsRNA in an ATP-dependent manner (Huang & Liu, 2002; Ishizuka *et al*, 2002). Rm62 recognizes stem loop structures and was previously shown to facilitate miRNA processing and antiviral defense against the Rift Valley fever virus (RVFV) in drosophila both *ex vivo* and *in vivo* (Moy *et al*, 2014). Therefore, our results could suggest that Rm62 may be recruited to a stem loop structure of the DCV RNA to facilitate its replication.

The protein Rin belongs to the GTPase activating protein (SH3 domain) binding protein family (also known as G3BPs) a family of RBPs that regulate gene expression in response to environmental stresses. In unstressed cells, Rin plays a role in the stabilization of target mRNAs and upregulation of their translation (Laver *et al*, 2020). This regulation can happen inside or outside stress granules (SGs), of which Rin is a core component, or outside of them (Alam & Kennedy, 2019). As SGs are induced in response to viral infections, some viruses sequestrate G3BPs (Lloyd, 2013) while others such as the poliovirus (*Picornaviridae* family) target and cleave them in order to disrupt the formation of stress granules (White *et al*, 2007). This is interesting as DCV is a picorna-like virus. In some cases, e.g. in alphaviruses, a conserved interaction with the viral proteins Nsp3 and SG proteins (Nowee *et al*, 2021) not only prevents SG formation but also act in favor of the virus, as in the case of the Rin-Nsp3 interaction for the chikungunya (CHIKV) virus (Fros *et al*, 2015). CHIKV is able to utilize Rin in order to increase infection rate and transmissibility. Interestingly, although we saw an antiviral effect of Rin and a proviral effect of Lig during DCV infection, the two proteins have been shown to act in concert to regulate cell proliferation during development (Baumgartner *et al*, 2013). In addition, the human homologue of Lig, UBAP2L, was identified after pull-down of the human homologue of Rin, G3BP1, in HEK293T cells (Markmiller *et al*, 2018). A possible explanation to these opposite effects could be that the disruption of this interaction in Lig KD flies allows Rin to focus on its antiviral function, and the absence of Lig would therefore be detrimental to the virus.

In conclusion, by using the combination of a global analysis with line-specific analyses, we were able to both obtain a global overview of the RNP network surrounding Dicer-2 and focus on the impact of specific factors on this network, such as the impact of the infection and different point mutations. Overall, this work has produced a resource and a large amount of data that is now available for the community. Our network hints at a potential involvement of Dicer-2 and several candidates in different mechanisms, in both mRNA regulation and antiviral immunity.

## MATERIAL AND METHODS

### Flies, drosophila S2 cells and virus

The GFP::Dicer-2 fusion drosophila fly lines were obtained by crossing [*w^IR^*; *dicer-2^L811fsx^*/*CyO*] virgins with: [*w^IR^*; *dicer-2^L811fsx^*/*CyO*; ubi>GFP::Dicer-2^WT^] males (GFP::Dicer-2^WT^), [GFP::Dicer-2^Hel1^ : *w^IR^*; *dicer-2^L811fsx^*/*CyO*; ubi>GFP::Dicer-2^G31R^] males (GFP::Dicer-2^Hel1^), or [*w^IR^*; *dicer-2^L811fsx^*/*CyO*; ubi>GFP::Dicer-2^E1371K/E1617K^] males (GFP::Dicer-2^RNaseIII^). The first two fly lines (GFP::Dicer-2^WT^ and GFP::Dicer-2^Hel1^), were established as described previously (Donelick *et al*, 2020) and the third one, GFP::Dicer-2^RNaseIII^, was established in the same manner with the following mutations in the Dicer-2 sequence: E1371K and E1617K, thus inactivating the activity of its RNase III domain. The control fly lines used were *CantonS* flies and GFP expressing flies, obtained by crossing *P{UAS-GFP.S65T}Myo31DF[T2]* line (BDSC #1521) under the control of the *actin5C* promoter (BDSC #25374). All experiments were performed with an equivalent number of males and females.

S2 cells were grown in Schneider’s medium (Biowest) supplemented with 10% fetal calf serum, Glutamax (Invitrogen) and Penicillin/Streptomycin (100x mix, 10 mg/mL/ 10000 U, Invitrogen). DCV virus stock was produced as described (Kemp *et al*, 2013).

### Viral infection and immunoprecipitations for MS analysis

Infections were performed on 3- to 5-day-old flies. Forty flies (20 females and 20 males) of each phenotype were injected with TRIS (10 mM, pH7.5) or DCV (500PFU) with 4.6 nL of TRIS (10 mM pH7.5) or DCV (500PFU) by intrathoracic injection (Nanoject II apparatus; Drummond Scientific). Flies were kept at 25°C for either 2 or 3 days, and then collected in Precellys tubes with zirconium beads and frozen overnight at -80°C. The flies were first shredded at 10°C without any buffer and then a second time with 1 mL lysis buffer (30 mM HEPES KOH pH 7.5, 50 mM NaCl, 2 mM Mg(OAc)2, 1% NP40, 2X cOmplete Protease Inhibitor Cocktail [Roche]). If the impact of RNase A on the interactants was tested, samples were separated in two halves and one half was treated with 15 µg RNase A for 30 min at 4°C. Samples were then centrifuged and the rest of the protocol was performed in the same manner as for the other MS experiment.

After centrifugation, 20 µL (0.05%) of the protein supernatant was kept to be loaded on a gel later on (**Supplementary Figure S1A**) and the rest was incubated with anti-GFP bead (Miltenyi) for 40 min at 4°C on a spinning wheel. The protein samples were then added onto a microcolumn placed on an uMACS separator (Miltenyi) after equilibration of the microcolumn with lysis buffer. Three washes were performed using wash buffer (30 mM HEPES KOH pH 7.5, 50 mM NaCl, 2 mM Mg(OAc)2, 0.1% NP40, 2X cOmplete Protease Inhibitor Cocktail [Roche]) and the elution was performed using elution buffer (Miltenyi) heated to 95°C. Five microliters (25%) of the elution was kept to confirm the IP on a WB and the rest was immediately used for mass spectrometry analysis.

### Mass spectrometry analysis

Proteins were digested with sequencing-grade trypsin (Promega, Fitchburg, MA, USA). Each sample was analyzed by nanoLC-MS/MS on a QExactive+ mass spectrometer coupled to an EASY-nanoLC-1000 (Thermo-Fisher Scientific, USA) as described previously (Chicher et al., 2015). Data were searched against the Drosophila melanogaster UniprotKB sub-database with a decoy strategy (UniprotKB release 2018_02 and 2022_01, taxon 7227, 42551 forward protein sequences). The correlation between spectrum matches for common baits of the two different MS datasets was assessed, as the two MS experiments were performed using different UniprotKB releases, and was greater than 0.999. Peptides and proteins were identified with Mascot algorithm (version 2.5.1, Matrix Science, London, UK) and data were further imported into Proline v1.4 software (http://proline.profiproteomics.fr/). Proteins were validated on Mascot pretty rank equal to 1, and 1% FDR on both peptide spectrum matches (PSM score) and protein sets (Protein Set score). The total number of MS/MS fragmentation spectra was used to quantify each protein from at least three independent biological replicates. The mass spectrometric data were deposited to the ProteomeXchange Consortium *via* the PRIDE partner repository with the dataset identifier PXD038898 and 10.6019/PXD038898.

### Statistical post-processing of the MS data and bioinformatics analysis

For each fly line, statistical post-processing of the data was performed through R (v3.2.5), using the IPinquiry4 package (Kuhn *et al*, 2023). After a column-wise normalization of the data matrix, the spectral count values were submitted to a negative-binomial test using an edgeR GLM regression as well as the msmsTests R package (release 3.6, Gregori et al., 2013). For each identified protein, an adjusted *p*-value (adjp) corrected by Benjamini–Hochberg was calculated, as well as a protein fold-change (FC). The results are presented in a Volcano plot using protein log2 fold changes and their corresponding adjusted log10 *p*-values to highlight upregulated proteins. Duplicate genes were manually removed and the results of the lists of genes enriched in the different conditions were put in JVenn to produce Venn diagrams (Bardou *et al*, 2014), or in R using the Pheatmap function to produce heatmaps.

An overall analysis of the data was performed using the Significance Analysis of INTeractome (SAINTexpress, v3.6.1) tool for interaction scoring. The data was then represented using Prohits-viz (primary filter 0.01, secondary filter 0.05, Ward clustering method and Camberra distance metric). A protein network was represented using the STRING tool in Cytoscape (version v3.9.1), after inputting the top 15% of the SAINT analysis candidates. The GO term functional enrichment analysis was conducted with STRING (https://string-db.org), using the full protein list and log fold-change.

### Confirmation of the interactions by immunoblot analysis

The input and elution samples obtained after immunoprecipitation using anti-GFP beads (Miltenyi or agarose beads from Chromotek). If necessary, injections followed by RNase A treatment or not were performed as described above. Protein extracts were separated on a Biorad gel and transferred to a nitrocellulose membrane. Membranes were then blocked in TBST containing 5% milk powder for 1h at RT and incubated overnight at 4°C with the different primary antibodies listed in **Supplementary Table 2**. After washing, the corresponding secondary antibodies fused to horseradish peroxidase (HRP) were added to the membrane for 2h at RT. Membranes were then washed and visualized with enhanced chemiluminescence reagent (GE Healthcare) in a ChemiDoc (Bio-Rad) apparatus.

### RNAi screen in S2 cells

After clustering of the 288 candidates highlighted by SAINT analysis after LC-MS/MS, the top 15% were selected to be part of a RNAi mini-screen. The dsRNAs were ordered at Drosophila RNAi Screening Center (DRSC) at Harvard Medical School. 47 dsRNAs were ordered in total, amongst which *dsAGO2* and *dsRACK1*, which were used as antiviral and proviral controls, respectively. Moreover, two negative controls (*dsGFP* and *dsLacZ*) and on RNAi knockdown (KD) efficiency control (*dsThread*) were placed in triplicate for each plate and put at different positions on the plate. *dsGFP* and *dsLacZ* were used to detect and normalize column and/or line effect, and *dsThread* controls were used to check that RNAi was working properly for each plate. Cells were seeded in 96-well plates and incubated with 1µg of dsRNA in FBS-free Schneider medium for 4h. After this starving period, normal S2 medium containing FBS was added, and the cells were incubated for 3 days before infection. Infections were performed at 25°C with MOI of 0.01 for 20h, after adsorption on ice for 1h. Cell lysis and reverse transcription were then performed using the Cells-to-CT kit (ThermoFisher Scientific) according to the manufacturer protocol, and used to perform quantitative real-time qPCR using the iTaq Universal SYBR Green Supermix (Biorad). The primers used are listed in the **Supplementary Table 3**. After calculating the 2^ΔCt^ for each sample, different mixed effect linear models were tested to search for bias in the data (lme2 R package (version 1.1-29)). As variations in plate, rows and columns had a significant impact after ANOVA, they were used as random factors in the final mixed effect model chosen. Corrected ratios (estimates) were extracted from the model and statistical significance was calculated by comparing estimates of each dsRNA to that of the dsLacZ control using the emmeans function.

### Genetic UAS-RNAi screen in an *actin5C*-Gal4;*tub*-Gal80^TS^ system

The top 15% candidates highlighted by the SAINT analysis were selected for the genetic screen *in vivo*. KK and GD inverted repeat transgenic fly lines for each candidate gene were acquired from the VDRC stock center (**Supplementary Table 4**), and *shmCherry* (BDSC #35787) and *shAGO2* (BDSC #34799) were used as controls, in addition to the respective KK and GD control lines. Transgenic males containing the inverted repeat of the target gene under the control of Gal4 regulated upstream activating sequence (UAS) were crossed with virgin females [*actin5*C-Gal4/*CyO*; *tubulin*-Gal80^TS^] at 18°C. The F1 generation was placed at 29°C for 5 days to induce the knockdown of candidate genes. All experiments were subsequently performed at 29°C. Infections were then performed as described above. The flies were then either counted every day to assess survival or collected at 1 or 2 dpi for RT-qPCR. In this case, three males and three females per condition were collected. Total RNA from the flies was then extracted using Trizol-chloroform, and 500 ng of total RNA was reverse transcribed using the iScript™ gDNA Clear cDNA Synthesis Kit (Biorad) according to the manufacturer’s instructions, and used to perform quantitative real-time qPCR using the iTaq Universal SYBR Green Supermix (Biorad), on a CFX384 Touch Real-Time PCR platform (Bio-Rad). The qPCR primers used are listed in **Supplementary Table 2**.

### Normalization and qPCR/survival analysis

All statistical analyses were performed in R (version 3.6.1). To perform the qPCR analysis, after calculating the 2^ΔCt^ for each sample, different mixed effect linear models were tested to search for bias in the data (lme2 R package (version 1.1-29)). Each batch was analyzed independently, and the data was tested for normality and homoscedasticity. As variations in replicates had a significant impact after ANOVA, this parameter was used as a random factor in the final mixed effect model chosen. Moreover, dependence between the “Gene” and “dpi” fixed factors was tested and found to be true for Batch 3; the models were adjusted accordingly. Corrected ratios (estimates) were extracted from the model and statistical significance was calculated by comparing estimates of each fly line to that of the shmCherry control using the emmeans function. P-values were adjusted using the Dunnett method. For survival analysis, the hazard ratio for each candidate was computed using a Cox Proportional-Hazards Model. In addition, Kaplan-Meier curves for each gene were produced using the Survival package, and the corresponding logrank tests were performed to compare each candidate to shmCherry.

## Acknowledgments

This work of the Interdisciplinary Thematic Institute IMCBio, as part of the ITI 2021-2028 program of the University of Strasbourg, CNRS and Inserm, was supported by IdEx Unistra (ANR-10-IDEX-0002), and by SFRI-STRAT’US project (ANR 20-SFRI-0012) and EUR IMCBio (ANR-17-EURE-0023) under the framework of the French Investments for the Future Program. Part of this work was also funded by the ViroMOD project, under the framework of the Regional Cooperation Fund for Research (FRCR). The mass spectrometry instrumentation was funded by the University of Strasbourg IdEx “Equipement mi-lourd” 2015. The Rump monoclonal antibody developed by Gavis, E.R. was obtained from the Developmental Studies Hybridoma Bank, created by the NICHD of the NIH and maintained at The University of Iowa, Department of Biology, Iowa City, IA 52242.

## Author contribution

Claire Rousseau and Carine Meignin wrote the manuscript and designed the experiments. Claire Rousseau, Carine Meignin and Emilie Lauret performed the experiments. The mass spectrometry was performed by Lauriane Kuhn, Johana Chicher and Philippe Hammann and analysed by Claire Rousseau and Lauriane Kuhn. The rest of the analyses were performed by Claire Rousseau. We thank Jean-Luc Imler and Joao Marques for the critical reading of the manuscript, and Alfredo Castello for his scientific advice.

