## Supplementary figures for "Identification of new proviral and antiviral factors through the study of the Dicer-2 interactome *in vivo* during viral infection in *Drosophila melanogaster*"

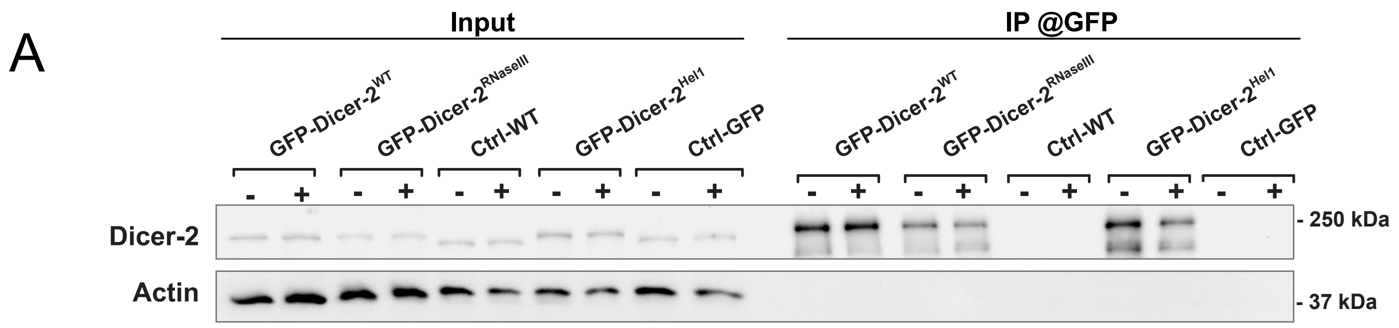


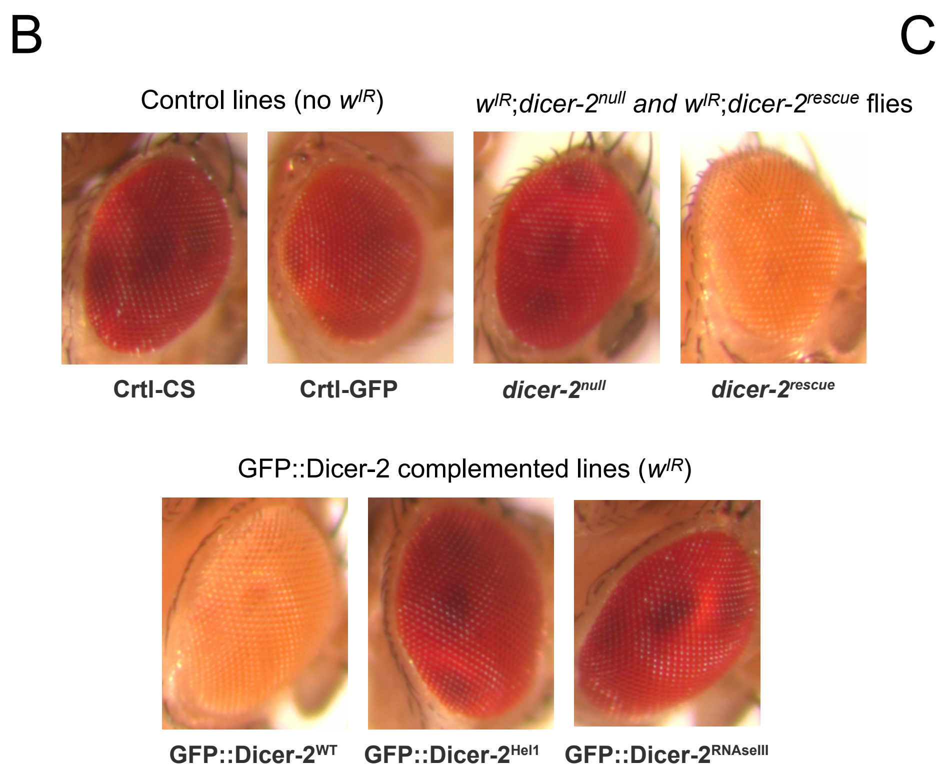


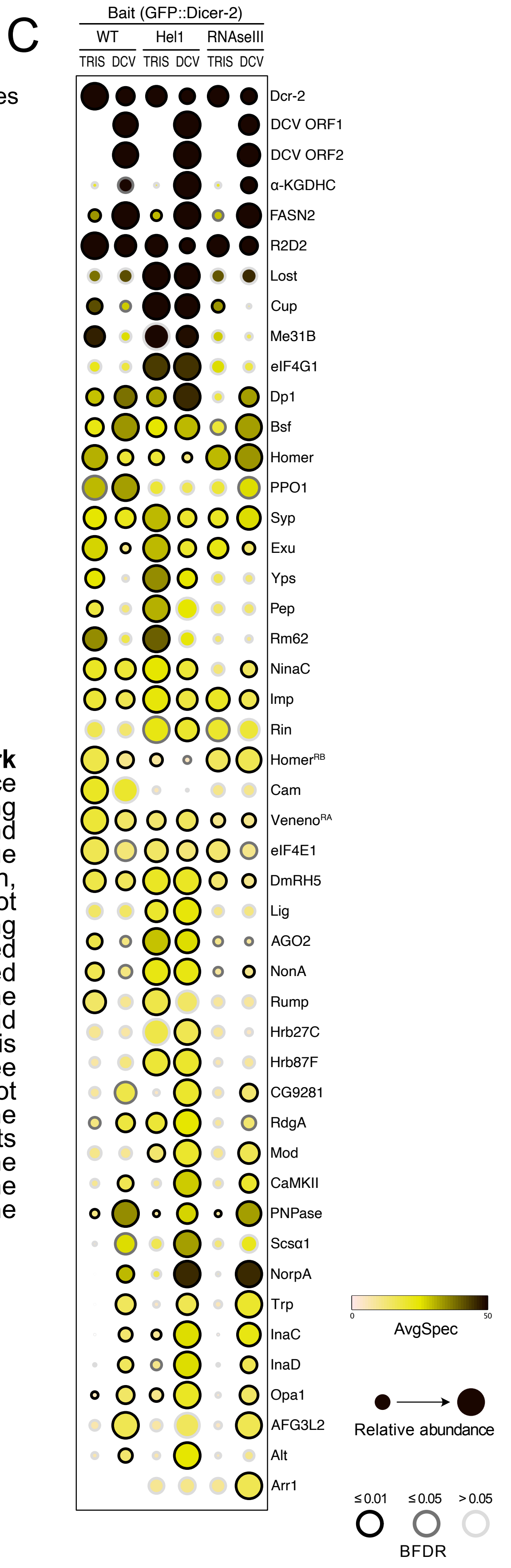


**Supplementary figure S1: Global Dicer-2 interactome network during viral infection *in vivo*. (A)** Immunoblot showing the presence of the Dicer-2 in all the samples before immunoprecipitation (IP) using anti-GFP beads, and in the GFP::Dicer-2 samples after IP. The band for the GFP::Dicer-2 fusion protein is slightly higher than Dicer-2 due to the added size molecular weight of the GFP, as expected. Actin, which does not interact with Dicer-2, can be seen in the input but not in the elution. **(B)** The color of the eyes of wIR flies allow the monitoring of RNAi efficiency. The *dicer-2*$^{null}$ and *dicer-2*$^{rescue}$ flies are described in Kemp et al., 2013. Flies without the transgene normally have red eyes (see Ctrl-CS and Ctrl-GFP). When the flies contain the transgene, if RNAi is efficient it induces KD of the *white* gene and results in a white eye phenotype (see *dicer-2*$^{rescue}$) and if RNAi is inefficient the KD is not effective and eyes are red like WT flies (see *dicer-2*$^{null}$). RNAi works normally in GFP::Dicer-2$^{WT}$, but not GFP::Dicer-2$^{Hel1}$ or GFP::Dicer-2$^{RNaseIII}$ flies. **(C)** Top 15% of the candidates highlighted by the IP-MS experiment. The color represents the average number of spectra (AvgSpec) detected in the samples, the size of the dot represent the relative abundance of the protein in the condition compared to the controls, and the color of the outline of the dot represents the BFDR (Baysian False Discovery Rate).

| Ranking | Gene name | Accession number |
|---|---|---|
| Bait | Dicer-2 | A1ZAW0 |
| Bait | GFP::Dcr-2$^{WT}$ | - |
| Bait | GFP::Dcr-2$^{RNase III}$ | - |
| Bait | GFP::Dcr-2$^{Hel1}$ | - |
| Virus | ORF1 | O36966 |
| Virus | O36967 | O36967 |
| 1 | alpha-KGDHC | Q9VGQ1 |
| 2 | FASN2 | M9PB21 |
| 3 | R2D2 | Q2Q0K7 |
| 4 | lost | Q9VN21 |
| 5 | cup | Q9VMA3 |
| 6 | Me31B | P23128 |
| 7 | eIF4G1 | A8DZ29 |
| 8 | Dp1 | Q7KN75 |
| 9 | bsf | Q95NR4 |
| 10 | homer | O96607 |
| 11 | PPO1 | Q27598 |
| 12 | Syp | A0A0B4KGF9 |
| 13 | exu | P28750 |
| 14 | yps | Q95RE4 |
| 15 | Pep | M9NG39 |
| 16 | Rm62 | P19109 |
| 17 | ninaC | P10676 |
| 18 | Imp | M9NF14 |
| 19 | rin | Q9NH72 |
| 20 | homer-RB | E8NH56 |
| 21 | Cam | P62152 |
| 22 | Veneno | N0A2N3 |
| 23 | eIF4E1 | P48598 |
| 24 | CG10077 | Q8MZI3 |
| 25 | lig | Q86S05 |
| 26 | AGO2 | Q9VUQ5 |
| 27 | nonA | Q04047 |
| 28 | rump | Q9VHC7 |
| 29 | Hrb27C | P48809 |
| 30 | Hrb87F | P48810 |
| 31 | CG9281 | Q9VXR5 |
| 32 | rdgA | A0A023GPM5 |
| 33 | mod | P13469 |
| 34 | CaMKII | A4V134 |
| 35 | PNPase | Q5U1D1 |
| 36 | Scsα1 | Q94522 |
| 37 | norpA | P13217 |
| 38 | trp | P19334 |
| 39 | inaC | P13677 |
| 40 | inaD | Q24008 |
| 41 | Opa1 | A0A0B4LGF5 |
| 42 | AFG3L2 | Q8T4G5 |
| 43 | alt | Q960Y8 |
| 44 | Arr1 | P15372 |
| 45 | CG1814 | A1Z7V9 |
| 46 | Acp26Aa | P10333 |
| 47 | Arr2 | P19107 |
| 48 | CG10737 | B7YZL1 |
| 49 | jp | Q7JV09 |
| 50 | Lsd-1 | A0A0B4KHZ1 |
| 51 | eIF3m | Q7JVI3 |
| 52 | AspRS | Q7K0E6 |
| 53 | alphaCOP | Q9W0B8 |
| 54 | Dhc64C | P37276 |
| 55 | eIF3i | O02195 |
| 56 | eIF3d1 | Q9VCK0 |
| 57 | eEF1gamma | Q9NJH0 |
| 58 | eIF3c | A1ZAX1 |
| 59 | GluProRS | P28668 |
| 60 | Cyp4g1 | Q9V3S0 |
| 61 | kdn | Q9W401 |
| 62 | ORF | Q23992 |
| 63 | Hsp26 | P02517 |
| 64 | lva | Q8MSS1 |
| 65 | E3 | Q9VVL7 |
| 66 | Map205 | P23226 |
| 67 | Hsp27 | P02518 |
| 68 | AGO1 | Q32KD4 |
| 69 | larp-RB | F9W325 |
| 70 | Arc1 | Q7K1U0 |
| 71 | Zn72D | Q86BI3 |
| 72 | loqs | Q9VJY9 |
| 73 | eIF4A | Q02748 |
| 74 | mEFTu2 | Q7K3V6 |
| 75 | clu | A1ZAB5 |
| 76 | mahe | Q9W3M7 |
| 77 | Hsc70-5 | P29845 |
| 78 | FK506-bp1 | P54397 |
| 79 | par-1 | E1JGN0 |
| 80 | EP(2)2054 | A1Z9K0 |
| 81 | piwi | Q9VKM1 |
| 82 | 128up | P32234 |
| 83 | Tailor | A0A0B4KGN4 |
| 84 | Hem | P55162 |
| 85 | Cdep | A0A0C4DHA1 |
| 86 | ps | Q0KI96 |
| 87 | Sra-1 | Q9VF87 |
| 88 | CG15784 | A9YIM2 |
| 89 | CG5641 | Q9VG73 |
| 90 | lark | Q94901 |
| 91 | RtcB | Q9VIW7 |
| 92 | gag-r | P91787 |
| 93 | RnpS1 | Q9VHC0 |
| 94 | Cbp80 | Q7K4N3 |
| 95 | Ge-1 | Q9VKK1 |
| 96 | rig | Q86BY9 |
| 97 | ND-B14.5B | Q9VQM2 |
| 98 | eIF4G2 | Q9VCH1 |
| 99 | Ostgamma | Q8SY53 |
| 100 | fest | A0A0B4JCT9 |
| 101 | TrpRS-m | Q8SZU2 |
| 102 | GIP | P04146 |
| 103 | Adar | M9NEQ7 |
| 104 | CG17593 | Q9VQR9 |
| 105 | CG5913 | Q960C1 |
| 106 | Not1 | A0A0B4LEZ3 |
| 107 | BEST:CK01174 | Q9VSH0 |
| 108 | Nsf2 | P54351 |
| 109 | Lon | Q7KUT2 |
| 110 | CG10912 | E1UI89 |
| 111 | DIP1 | A4V4V2 |
| 112 | gammaCOP | A0A0B4KHL2 |
| 113 | FASN3 | Q7PLB8 |
| 114 | ctp | Q24117 |
| 115 | Acsl | A0A0B4KFE4 |
| 116 | NO66 | Q7K4H4 |
| 117 | eIF3b | Q0E940 |
| 118 | Galk | Q95U34 |
| 119 | MetRS | A1ZBE9 |
| 120 | CG16935 | Q9V6U9 |
| 121 | Tango5 | Q9W2S1 |
| 122 | CG8360 | Q9VLR3 |
| 123 | AP-1-2beta | Q24253 |
| 124 | CG13850 | Q9VD14 |
| 125 | Hexo2 | Q9W3C4 |
| 126 | TM4SF | A0A0B4KFZ4 |
| 127 | CG6255 | A0AMM0 |
| 128 | Echs1 | Q7JR58 |
| 129 | vari | M9PDB2 |
| 130 | Lmpt | M9MRX0 |
| 131 | r | P05990 |
| 132 | CCT4 | Q9VK69 |
| 133 | Klp10A | Q960Z0 |
| 134 | rept | Q9V3K3 |
| 135 | shv | Q9VPQ2 |
| 136 | wal | Q7KLW5 |
| 137 | Sac1 | Q9W0I6 |
| 138 | eIF2alpha | P41374 |
| 139 | CG8768 | E1UIK3 |
| 140 | BcDNA:GH07921 | Q9VT61 |
| 141 | gag | P10405 |
| 142 | sm | A0A0B4K7B0 |
| 143 | Larp4B | Q9I7T7 |
| 144 | CG5787 | Q9VK59 |
| 145 | Stip1 | Q9VPN5 |
| 146 | beg | Q8T3L6 |
| 147 | scu | O18404 |
| 148 | Amph | A0A0B4KEW6 |
| 149 | pont | Q9VH07 |
| 150 | shrb | Q8T0Q4 |
| 151 | Kank | A0A0B4JCT6 |
| 152 | Glg1 | Q9VP27 |
| 153 | BcDNA:RE30174 | Q9VZZ6 |
| 154 | yl | P98163 |
| 155 | CG3967 | Q9VSY4 |
| 156 | Hsp22 | P02515 |
| 157 | Trap1 | A1Z6L9 |
| 158 | Top3beta | O96651 |
| 159 | CCT7 | Q9VHL2 |
| 160 | SRPK | A0A0B4KF69 |
| 161 | Smn | Q9VV74 |
| 162 | rtp | Q9VN91 |
| 163 | qkr58E-1 | A0A0B4LG88 |
| 164 | Ddx1 | Q9VNV3 |
| 165 | CG31879 | Q7KTG0 |
| 166 | Q6AWL1 | Q6AWL1 |
| 167 | DCP1 | Q9W1H5 |
| 168 | CG3071 | O46069 |
| 169 | san | Q9NHD5 |
| 170 | Edc3 | Q9VVI2 |
| 171 | Psi | A1ZAK7 |
| 172 | Upf1 | Q9VYS3 |
| 173 | CG2246 | A0A0B4JD23 |
| 174 | DnaJ-1 | Q24133 |
| 175 | rgn | Q9Y102 |
| 176 | His3; | P02299 |
| 177 | dhd | P47938 |
| 178 | nocte | M9PE74 |
| 179 | Atx2 | Q8SWR8 |
| 180 | FeCh | Q9V9S8 |
| 181 | Pabp2 | Q7KNF2 |
| 182 | Abi | A0A0B4K774 |
| 183 | Iswi | Q24368 |
| 184 | Pss | M9PIC3 |
| 185 | AspRS-m | Q9VJH2 |
| 186 | Sf3b3 | Q5BI86 |
| 187 | mask | Q9VCA8 |
| 188 | Cand1 | Q9VKY2 |
| 189 | Mccc1 | Q9V9T5 |
| 190 | Cpr49Ab | A1Z8Y3 |
| 191 | Tdrd3 | Q9VUH8 |
| 192 | l(1)G0020 | Q9W3C1 |
| 193 | tyf | E1JJD6 |
| 194 | hnRNP | A1ZBB4 |
| 195 | stau | P25159 |
| 196 | Q9VU21 | Q9VU21 |
| 197 | mub | A4IJ59 |
| 198 | CG5728 | Q9VC94 |
| 199 | mtgo | B7YZW3 |
| 200 | mle | P24785 |
| 201 | SCAR-RA | C1C3C5 |
| 202 | aub | O76922 |
| 203 | Pp1alpha-96A | P48461 |
| 204 | CG7382 | Q9VMR0 |
| 205 | numb | P16554 |
| 206 | Sec22 | O77434 |
| 207 | Q494I6 | Q494I6 |
| 208 | Drp1 | Q8IHG0 |
| 209 | RanBPM | Q4Z8K6 |
| 210 | Caf1-55 | Q24572 |
| 211 | MEP-1 | Q0E8J0 |
| 212 | stck | Q8IGP6 |
| 213 | AIMP3 | Q8MKK1 |
| 214 | Myo31DF | Q23978 |
| 215 | LeuRS | Q8MRF8 |
| 216 | CCT6 | Q9VXQ5 |
| 217 | CG14445 | A8JUZ6 |
| 218 | CG44242 | B7YZH7 |
| 219 | Vha36-1 | Q9V7D2 |
| 220 | gish | A0A0B4K697 |
| 221 | gammaTub37C | P42271 |
| 222 | woc | A0A0B4KHZ0 |
| 223 | epsilonCOP | Q9Y0Y5 |
| 224 | ND-B17.2 | Q8MSI7 |
| 225 | RnrL | P48591 |
| 226 | NAT1 | A1Z968 |
| 227 | Rpn13 | Q7K2G1 |
| 228 | Ns1 | Q8MT06 |
| 229 | tud | P25823 |
| 230 | Tor | Q9VK45 |
| 231 | Marf | Q7YU24 |
| 232 | EG:115C2.2 | O77426 |
| 233 | CkIalpha | P54367 |
| 234 | caz | Q27294 |
| 235 | Dcr-1 | Q9VCU9 |
| 236 | AcCoAS | Q9VP61 |
| 237 | Sarm | Q6IDD9 |
| 238 | eRF1 | Q9VPH7 |
| 239 | IM33 | Q9VQT8 |
| 240 | CCT1 | P12613 |
| 241 | HIP | C4NYP8 |
| 242 | eIF2Bepsilon | Q9W541 |
| 243 | Cisd2 | Q9VAM6 |
| 244 | Vps2 | Q9VBI3 |
| 245 | Tctp | Q9VGS2 |
| 246 | Ist1 | M9PEC1 |
| 247 | CG6178 | Q9VCC6 |
| 248 | Pi4KIIIalpha | M9PDM4 |
| 249 | ArgRS | Q9VXN4 |
| 250 | BEST:GH19547 | A1ZAU4 |
| 251 | BEST:GH10831 | Q6NMY2 |
| 252 | IleRS | Q8MSW0 |
| 253 | nwk | M9NE66 |
| 254 | tacc | A0A0B4JCV4 |
| 255 | Pkc53E | P05130 |
| 256 | Syt7 | H9XVN2 |
| 257 | eIF2beta | P41375 |
| 258 | msn | Q9W002 |
| 259 | Sgs7 | P02841 |
| 260 | ens | M9PE93 |
| 261 | Chro | Q8T9D1 |
| 262 | Prp31 | Q9VUM1 |
| 263 | fit | Q9VD66 |
| 264 | CG3756 | Q9VMX3 |
| 265 | OstDelta | Q7K110 |
| 266 | msps | A0A0B4K664 |
| 267 | ND-13B | Q9VTB4 |
| 268 | CG7300 | Q9VKR7 |
| 269 | sle | Q8INM3 |
| 270 | stnB | Q24212 |
| 271 | CG33523-RD | I0DHK9 |
| 272 | cg12493 | M9PHF9 |
| 273 | Q4QPU9 | Q4QPU9 |
| 274 | MRG15 | Q9Y0I1 |
| 275 | HnRNP-K | A1ZBW0 |
| 276 | BicC | Q24009 |
| 277 | peng | O61345 |
| 278 | mago | P49028 |
| 279 | Sf3a1 | Q9VEP9 |
| 280 | Bx42 | P39736 |
| 281 | CG12128 | A1Z830 |
| 282 | Nnp-1 | Q9VJZ7 |
| 283 | CG8915 | Q9VX63 |
| 284 | Nopp140 | M9PFZ1 |
| 285 | mRpL12 | Q2XYH0 |
| 286 | CRIF | Q9VP13 |
| 287 | Rfc37 | Q9VX15 |
| 288 | Muc12Ea | Q8IR52 |

**Supplementary Table 2: Global Dicer-2 interactome.** List if 288 proteins that were highlighted by the global analysis using the SAINTexpress tool. The baits are represented in green, two viral proteins are in orange, the top 10% candidates are in blue, and the top 15% are in purple.

| Target | Reference | Animal | Dilution WB |
|---|---|---|---|
| Mouse IgG | M365FK (Rockland) | Goat | 1:10,000 |
| Rabbit IgG | NA934 (Amersham) | Goat | 1:10,000 |
| Dicer-2 | ab4732 (Abcam) | Rabbit | 1:500 |
| GFP | A11122 (Invitrogen) | Rabbit | 1:2,000 |
| R2D2 | ab14750 (Abcam) | Rabbit | 1:1,000 |
| DCV VP2 | 3593 (IGBMC) | Rabbit | 1:1,000 |
| DCV RdRp#2 | homemade | Guinea pig | 1:2,000 |
| eIF4E1 | Kindly provided by Izzauralde lab | rabbit | 1:1,000 |
| Rump | AB_10571461 (DSHB) | mouse | 1:300 |
| Lost | AB_2618045 (DSHB) | mouse | 1:500 |
| Me31b | Kindly provided by Izzauralde lab | rabbit | 1:1,000 |
| Syp | homemade | guinea pig | 1:2,000 |

**Supplementary Table 3: Antibodies used**

| Gene | Sequence | Orientation |
|---|---|---|
| RP49 | GCCGCTTCAAGGGACAGTATCT | Forward |
| RP49 | AAACGCGGTTCTGCATGAG | Reverse |
| DCV | TCATCGGTATGCACATTGCT | Forward |
| DCV | CGCATAACCATGCTCTTCTG | Reverse |

**Supplementary Table 4: Primer used for qPCR**

| Gene | Reagent ID | dsRNA length (bp) |
|---|---|---|
| LacZ | Ctrl | 593 |
| GFP | Ctrl | 609 |
| Thread | Ctrl | 809 |
| AGO2 | DRSC31768 | 265 |
| AGO2 | DRSC31769 | 239 |
| RACK1 | DRSC23796 | 512 |
| RACK1 | DRSC30688 | 255 |
| CG5214 | DRSC25298 | 363 |
| FASN2 | DRSC00610 | 511 |
| r2d2 | DRSC29562 | 319 |
| lost | DRSC34368 | 368 |
| lost | DRSC34034 | 266 |
| Cup | DRSC24610 | 465 |
| Cup | DRSC35662 | 206 |
| Me31B | DRSC03569 | 501 |
| Me31B | DRSC38084 | 301 |
| eIF4G1 | DRSC17218 | 503 |
| eIF4G1 | DRSC29786 | 379 |
| Dp1 | DRSC07403 | 511 |
| Dp1 | DRSC35700 | 290 |
| bsf | DRSC42668 | 545 |
| bsf | DRSC02035 | 505 |
| homer | DRSC29162 | 348 |
| PPO1 | DRSC05929 | 511 |
| PPO1 | DRSC40857 | 404 |
| Syp | DRSC34987 | 267 |
| Syp | DRSC29126 | 525 |
| exu | DRSC07615 | 515 |
| exu | DRSC42214 | 215 |
| yps | DRSC25035 | 234 |
| yps | DRSC11411 | 473 |
| Pep | DRSC37986 | 367 |
| Pep | DRSC11252 | 297 |
| Rm62 | DRSC32537 | 203 |
| Rm62 | DRSC32538 | 159 |
| ninaC | DRSC37305 | 346 |
| ninaC | DRSC36608 | 317 |
| Imp | DRSC20255 | 510 |
| Imp | DRSC35798 | 347 |
| rin | DRSC33098 | 220 |

| Gene | Reagent ID | dsRNA length (bp) |
|---|---|---|
| rin | DRSC33099 | 178 |
| Cam | DRSC07354 | 225 |
| Cam | DRSC07353 | 205 |
| Veneno | DRSC16537 | 500 |
| eIF4E1 | DRSC32114 | 203 |
| eIF4E1 | DRSC32113 | 195 |
| CG10077 | DRSC24756 | 348 |
| lig | DRSC34699 | 205 |
| lig | DRSC38413 | 537 |
| nonA | DRSC27455 | 283 |
| nonA | DRSC41353 | 140 |
| rump | DRSC16480 | 520 |
| rump | DRSC42718 | 366 |
| Hrb27C | DRSC28323 | 325 |
| Hrb27C | DRSC32135 | 265 |
| Hrb87F | DRSC29600 | 417 |
| Hrb87F | DRSC35793 | 320 |
| CG9281 | DRSC24522 | 435 |
| rdgA | DRSC35980 | 405 |
| rdgA | DRSC39133 | 307 |
| mod | DRSC28409 | 310 |
| CaMKII | DRSC17214 | 312 |
| CaMKII | DRSC36643 | 310 |
| PNPase | DRSC14304 | 506 |
| Scsalpha1 | DRSC26856 | 313 |
| Scsalpha1 | DRSC41030 | 212 |
| norpA | DRSC40745 | 534 |
| norpA | DRSC18806 | 163 |
| trp | DRSC17088 | 575 |
| trp | DRSC39226 | 492 |
| inaC | DRSC36646 | 383 |
| inaC | DRSC31368 | 292 |
| inaD | DRSC27116 | 367 |
| Opa1 | DRSC28908 | 429 |
| Opa1 | DRSC07191 | 511 |
| CG6512 | DRSC34403 | 391 |
| CG6512 | DRSC29981 | 331 |
| alt | DRSC32685 | 207 |
| alt | DRSC32684 | 160 |
| Arr1 | DRSC01867 | 513 |
| Arr1 | DRSC33429 | 245 |

**Supplementary Table 5: dsRNAs ordered from DRSC**

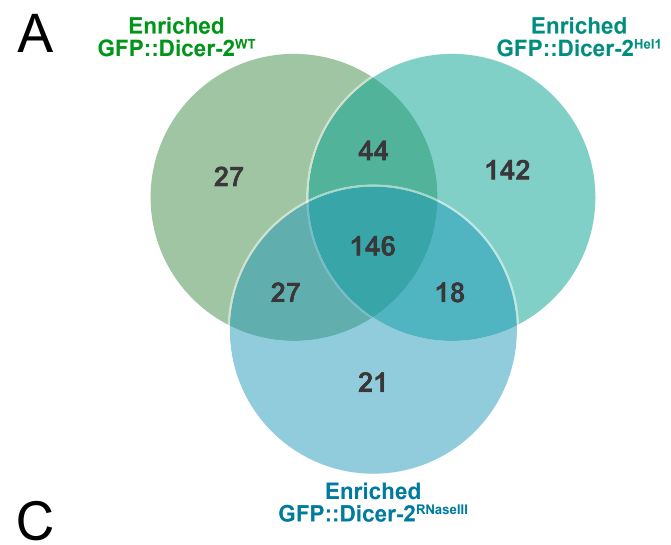


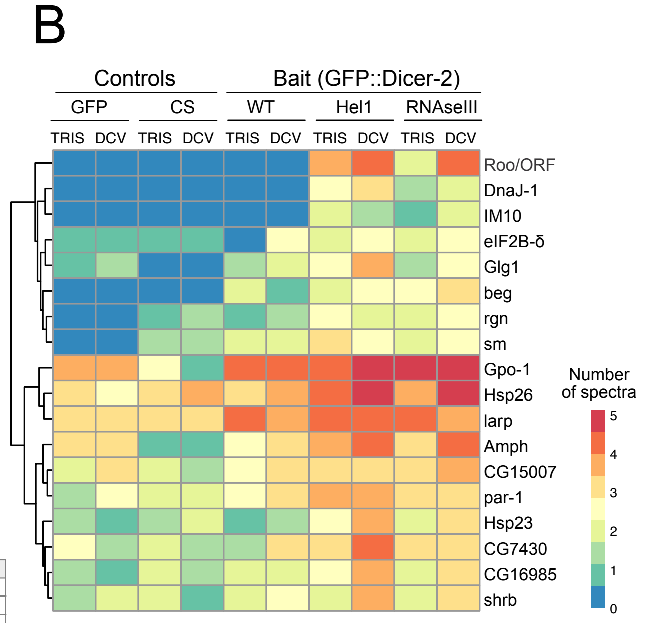


C

| Rank | Candidate | Rank | Candidate | Rank | Candidate | Rank | Candidate |
|---|---|---|---|---|---|---|---|
| 2 | FASN2 | 64 | lva | 124 | CG13850 | 193 | tyf |
| 3 | r2d2 | 66 | Map205 | 125 | Hexo2 | 195 | stau |
| 4 | lost | 67 | Hsp27 | 126 | TM4SF | 197 | mub |
| 5 | cup | 68 | AGO1 | 127 | CG6255 | 198 | CG5728 |
| 6 | Me31B | 69 | larp-RB | 129 | vari | 200 | mle |
| 8 | Dp1 | 70 | Arc1 | 130 | Lmpt | 201 | SCAR-RA |
| 9 | bsf | 71 | Zn72D | 131 | r | 202 | aub |
| 10 | homer | 72 | loqs | 133 | Klp10A | 203 | Pp1alpha-96A |
| 12 | Syp | 75 | clu | 134 | rept | 204 | CG7382 |
| 13 | exu | 77 | Hsc70-5 | 136 | wal | 205 | numb |
| 14 | yps | 78 | FK506-bp1 | 137 | Sac1 | 206 | Sec22 |
| 15 | Pep | 79 | par-1 | 139 | CG8768 | 208 | Drp1 |
| 16 | Rm62 | 81 | piwi | 141 | gag | 214 | Myo31DF |
| 18 | Imp | 82 | 128up | 142 | sm | 222 | woc |
| 19 | rin | 83 | Tailor | 144 | CG5787 | 224 | ND-B17.2 |
| 20 | homer-RB | 84 | Hem | 146 | beg | 225 | RnrL |
| 21 | Cam | 85 | Cdep | 147 | scu | 226 | NAT1 |
| 22 | Veneno-RA | 86 | ps | 148 | Amph | 228 | Ns1 |
| 23 | eIF-4E | 87 | Sra-1 | 149 | pont | 229 | tud |
| 25 | lig | 88 | CG15784 | 150 | shrb | 230 | Tor |
| 26 | AGO2 | 89 | CG5641 | 151 | Kank | 235 | Dcr-1 |
| 27 | nonA | 90 | lark | 152 | Glg1 | 239 | IM33 |
| 28 | rump | 92 | gag-r | 154 | yl | 241 | HIP |
| 29 | Hrb27C | 93 | RnpS1 | 158 | Top3beta | 251 | BEST:GH10831 |
| 30 | Hrb87F | 94 | Cbp80 | 160 | SRPK | 253 | nwk |
| 31 | CG9281 | 95 | Ge-1 | 161 | Smn | 254 | tacc |
| 32 | rdgA | 96 | rig | 162 | rtp | 256 | Syt7 |
| 33 | mod | 97 | ND-B14.5B | 163 | qkr58E-1 | 258 | msn |
| 34 | CaMKII | 98 | eIF4G2 | 164 | Ddx1 | 259 | Sgs7 |
| 37 | norpA | 99 | Ostgamma | 167 | DCP1 | 260 | ens |
| 38 | trp | 102 | GIP | 170 | Edc3 | 261 | Chro |
| 39 | inaC | 103 | Adar | 171 | Psi | 262 | Prp31 |
| 40 | inaD | 105 | CG5913 | 172 | Upf1 | 263 | fit |
| 41 | Opa1 | 106 | Not1 | 173 | CG2246 | 265 | OstDelta |
| 43 | alt | 108 | Nsf2 | 174 | DnaJ-1 | 266 | msps |
| 44 | Arr1 | 109 | Lon | 175 | rgn | 267 | ND-13B |
| 46 | Acp26Aa | 110 | CG10912 | 177 | dhd | 268 | CG7300 |
| 47 | Arr2 | 111 | DIP1 | 178 | nocte | 269 | sle |
| 48 | CG10737 | 112 | gammaCOP | 179 | Atx2 | 271 | CG33523-RD |
| 49 | jp | 113 | FASN3 | 180 | FeCh | 272 | cg12493 |
| 50 | Lsd-1 | 115 | Acsl | 182 | Abi | 275 | HnRNP-K |
| 53 | alphaCOP | 118 | Galk | 183 | Iswi | 276 | BicC |
| 54 | Dhc64C | 119 | MetRS | 187 | mask | 278 | mago |
| 60 | Cyp4g1 | 120 | CG16935 | 188 | Cand1 | 280 | Bx42 |
| 61 | kdn | 121 | Tango5 | 190 | Cpr49Ab | 281 | CG12128 |
| 62 | ORF | 122 | CG8360 | 191 | Tdrd3 | 282 | Nnp-1 |
| 63 | Hsp26 | 123 | AP-1-2beta | 192 | l(1)G0020 | 284 | Nopp140 |

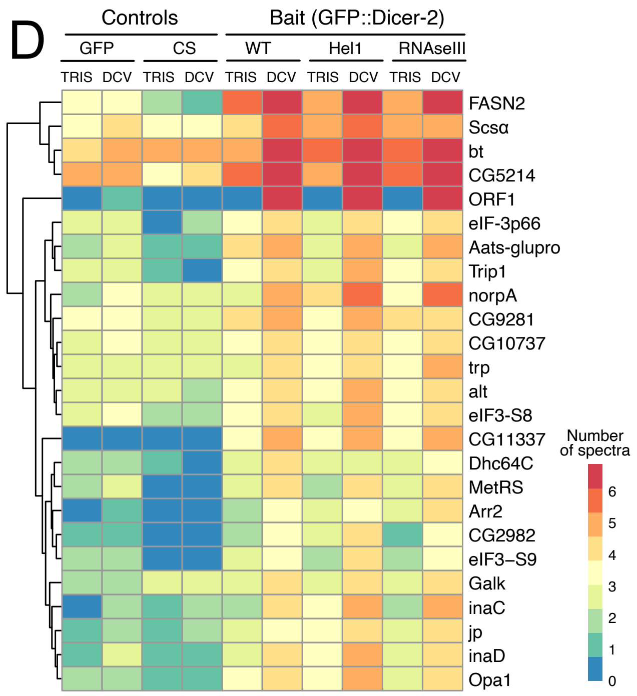


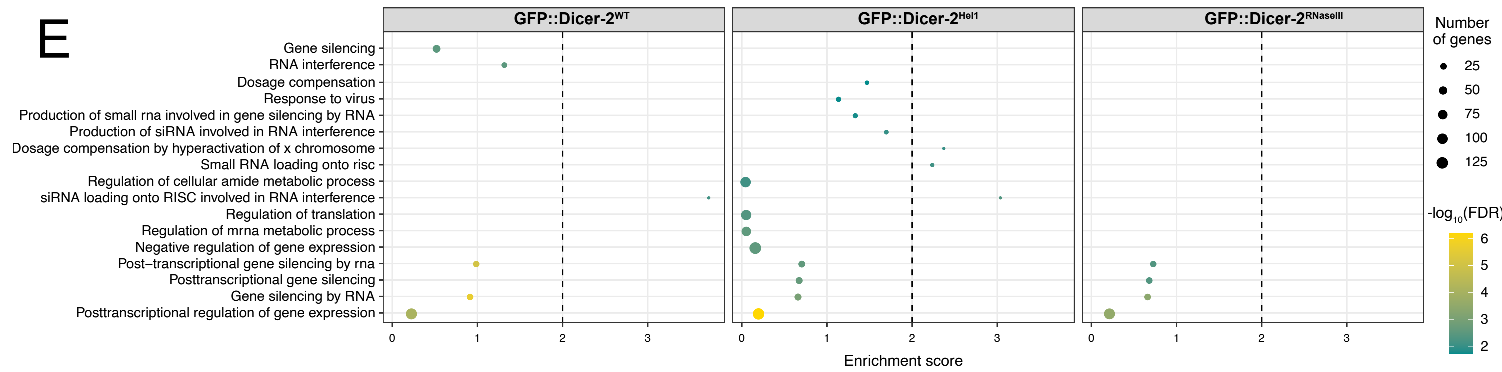


**Supplementary Figure S2: Impact of the Dicer-2 mutations on the RNP network of Dicer-2. (A)** Venn diagram showing the number of proteins identified in each GFP::Dicer-2 line. Candidates are selected with a fold-change > 2 and an adjusted *p*-value < 0.01. **(B)** Heatmap representing the number of spectra of the identified in each sample 18 proteins enriched only in the two GFP::Dicer-2 mutants (Hel1 and RNaseIII). (**C)** Table containing the candidates that were highlighted by both the global analysis and at least one of the line-specific analyses. **(D)** Heatmap representing the number of spectra of the 25 proteins enriched with all GFP::Dicer-2 lines during DCV infection. (E) GO term enrichment analysis "Biological Processes" for each GFP::Dicer-2 fly line compared to the controls.

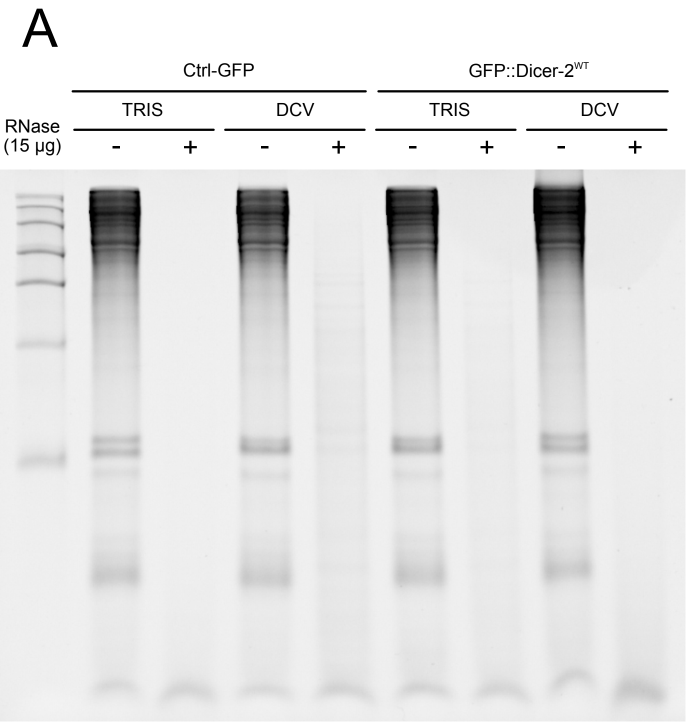


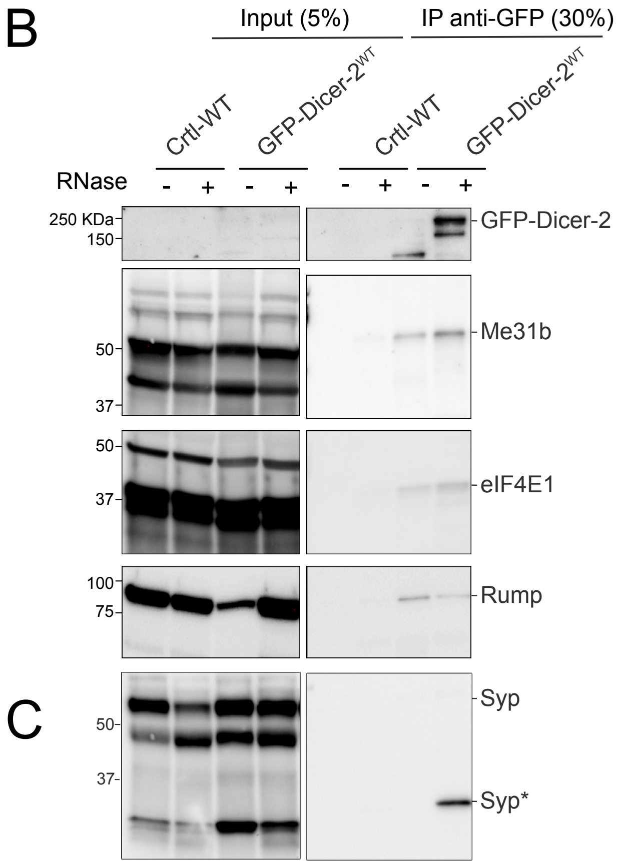


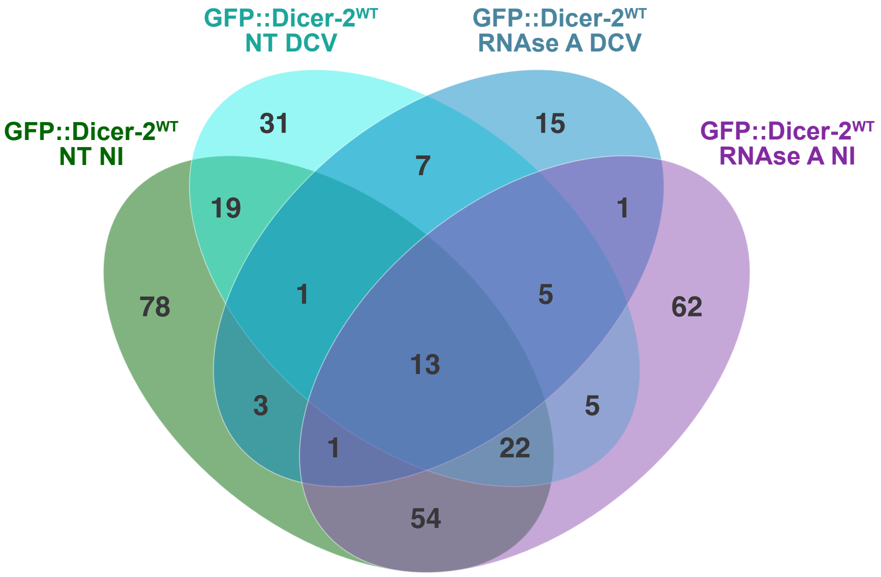


**Supplementary Figure S3: The RNA-dependent protein network of Dicer-2.** Urea/Acrylamide gel showing total RNA after no treatment or treatment with 15 µg of RNase A. Adult flies were injected or not with DCV and protein extraction was performed in the same manner as for the IPs. Instead of proceeding with the IP, total RNA was extracted after no treatment or treatment with RNase A to confirm the efficiency of the RNase treatment. **(B)** Venn diagram showing the number of proteins identified in the GFP::Dicer-2$^{WT}$ line in mock-infected and DCV-infected adult flies with or without RNAse A treatment. Candidates were selected with a fold-change > 2 and an adjusted *p*-value < 0.01. In total, 317 proteins were enriched in throughout the four different conditions, including some of the candidates from the global analysis. **(C)** Immunoprecipitation of GFP::Dicer-2$^{WT}$ in comparison to wilt-type CantonS flies (Ctrl-WT). GFP::Dicer-2$^{WT}$ interacts with Mei31B, eIF4E1, Rump and Syncrip.

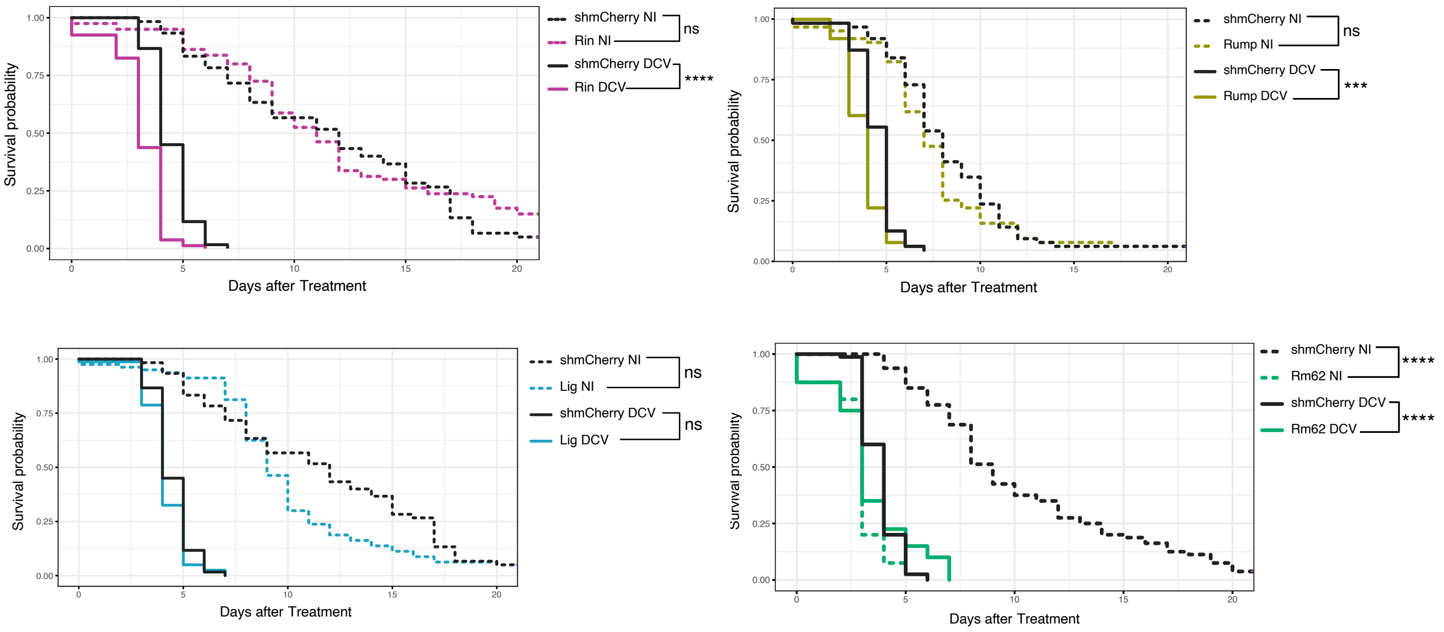


**Supplementary Figure S4: Role of the candidates in antiviral immunity in adult drosophila flies.** Survival analysis of KD Rin, Rump, Lig and RM62 *in vivo* after infection DCV 50 pfu. Significant survival compared to shmCherry line are represented in non infected (NI) and DCV infected.
